# TET2 regulates early and late transitions in exhausted CD8^+^ T-cell differentiation and limits CAR T-cell function

**DOI:** 10.1101/2024.03.29.587004

**Authors:** Alexander J. Dimitri, Amy E. Baxter, Gregory M. Chen, Caitlin R. Hopkins, Geoffrey T. Rouin, Hua Huang, Weimin Kong, Christopher H. Holliday, Volker Wiebking, Robert Bartoszek, Sydney Drury, Katherine Dalton, Owen M. Koucky, Zeyu Chen, Josephine R. Giles, In-Young Jung, Roddy O’Connor, Sierra Collins, John K. Everett, Kevin Amses, Scott Sherrill-Mix, Aditi Chandra, Naomi Goldman, Golnaz Vahedi, Julie K. Jadlowsky, Regina M. Young, Jan Joseph Melenhorst, Shannon L. Maude, Bruce L. Levine, Noelle V. Frey, Shelley L. Berger, Stephan A. Grupp, David L. Porter, Friederike Herbst, Matthew H. Porteus, Frederic D. Bushman, Evan W. Weber, E. John Wherry, Martha S. Jordan, Joseph A. Fraietta

**Author notes:** Authors contributed equally to this work (Alexander J. Dimitri, Amy E. Baxter, Gregory M. Chen, Martha S. Jordan, Joseph A. Fraietta).

## Abstract

CD8^+^ T-cell exhaustion hampers disease control in cancer and chronic infections and limits efficacy of T-cell−based therapies, such as CAR T-cells. Epigenetic reprogramming of CAR T-cells by targeting TET2, a methylcytosine dioxygenase that mediates active DNA demethylation, has shown therapeutic potential; however, the role of TET2 in exhausted T-cell (T_EX_) development is unclear. In CAR T-cell exhaustion models and chronic LCMV infection, TET2 drove the conversion from stem cell-like, self-renewing T_EX_ progenitors towards terminally differentiated and effector (T_EFF_)-like T_EX_. In mouse T-cells, *TET2*-deficient terminally differentiated T_EX_ retained aspects of T_EX_ progenitor biology, alongside decreased expression of the transcription factor TOX, suggesting that TET2 potentiates terminal exhaustion. TET2 also enforced a T_EFF_-like terminally differentiated CD8^+^ T-cell state in the early bifurcation between T_EFF_ and T_EX_, indicating a broad role for TET2 in mediating the acquisition of an effector biology program that could be exploited therapeutically. Finally, we developed a clinically actionable strategy for *TET2-* targeted CAR T-cells, using CRISPR/Cas9 editing and site-specific adeno-associated virus transduction to simultaneously knock-in a CAR at the *TRAC* locus and a functional safety switch within *TET2*. Disruption of *TET2* with this safety switch in CAR T-cells restrained terminal T_EX_ differentiation *in vitro* and enhanced anti-tumor responses *in vivo*. Thus, TET2 regulates pivotal fate transitions in T_EX_ differentiation and can be targeted with a safety mechanism in CAR T-cells for improved tumor control and risk mitigation.

**One Sentence Summary:** Modulation of exhausted CD8^+^ T-cell differentiation by targeting TET2 improves therapeutic potential of CAR T-cells in cancer.

## INTRODUCTION

T-cell exhaustion limits disease control in cancer and chronic viral infections. In the context of persistent antigen exposure, exhausted CD8^+^ T-cells (T_EX_) co-express multiple inhibitory receptors (IRs) and exhibit altered cytokine secretion, impaired proliferation and metabolic deficiencies compared to memory (T_MEM_) and effector (T_EFF_) T-cells (*1*). While T_EX_ cells hold significant clinical relevance, our understanding of T_EX_ development and the fundamental cellular and molecular mechanisms governing T_EX_ formation, maintenance, and activity, particularly in the setting of immunotherapies, is incomplete. Addressing these knowledge gaps could offer new strategies for enhancing patient outcomes.

Cellular immunotherapies such as chimeric antigen receptor (CAR) T-cells have transformed the treatment of cancer. However, T-cell exhaustion compromises the persistence and anti-tumor effector function of CAR T-cells *in vivo*, resulting in relapses in hematological malignancies and limited efficacy against solid tumors (*2*). Strategies including genetic modification of CAR T-cells to avert exhaustion (*3–6*) or use of immunotherapies such as PD1 blockade to reinvigorate T_EX_ cells have been proposed to enhance treatment efficacy (*7*). However, current reinvigoration strategies are insufficient to permanently reverse exhaustion (*8*), limiting therapeutic potential.

T_EX_ are a distinct epigenetic lineage and the discovery of regulators that govern the epigenetic remodeling events underpinning T_EX_ development holds promise for improving cell-based immunotherapies (*9, 10*). We previously reported massive clonal expansion of a single CAR T-cell in a patient undergoing therapy for chronic lymphocytic leukemia (CLL) that resulted in enhanced anti-tumor activity and subsequent complete and sustained disease remission (*10*). This unique clone had the CAR transgene integrated into *TET2*, accompanied by a pre-existing hypomorphic mutation in the patient’s second *TET2* allele. As a methylcytosine dioxygenase, TET2 plays a pivotal role in active DNA demethylation by initiating the conversion of 5-methylcytosine (5-mC) into 5-hydroxymethylcytosine (5-hmC), a first step in removal of the methyl group (*11, 12*). This finding suggested that modulation of TET2 could be used to alter the epigenetic landscape of T_EX_ and CAR T-cells for therapeutic benefit. Accordingly, biallelic disruption of *TET2*, with concomitantly sustained expression of the AP-1 factor BATF3, resulted in clonal proliferation of CAR T-cells with altered effector function (*13*). These data are consistent with skewed T_MEM_ versus T_EFF_ differentiation of *TET2*-deficient T-cells observed following acute viral infection (*14*). However, the role of TET2 in the precise T-cell fate transitions that govern T_EX_ cell differentiation remain unknown. Given that TET2 loss augments CAR T-cell efficacy (*10, ^13^*), unravelling the diverse roles of TET2 in the differentiation of CD8^+^ T-cells, particularly the epigenetic programming of T_EX_ cells across key developmental checkpoints, is critical for deciphering the underlying mechanisms of T-cell exhaustion and further enhancing the effectiveness of immunotherapies.

## RESULTS

### *TET2* is a frequent locus of transgene integration in CAR T-cell treated patients

Our previous report on *TET2*-disruption driving enhanced proliferation and sustained tumor clearance mediated by a single CAR T-cell in one patient (*10*) led us to investigate other potential occurrences of lentiviral integration into *TET2* in additional patients who underwent CAR T-cell therapy for CLL and acute lymphocytic leukemia (ALL) (*3*) (**tables S1 and S2**). Within these cohorts, 36% of CLL patients and 51% of ALL patients had at least one instance of lentiviral integration into *TET2* (**fig. S1A**). In total, 33 and 75 unique sites of integration were identified within CLL and ALL cohorts, respectively (**fig. S1B**). Most of these integration sites were of low abundance (**fig. S1C**) and occurred only once (**tables S1 and S2**); however, one CLL patient (p04409-09) exhibited a CAR transgene insertion at chr4+105190185 which was observed at multiple time points, at two weeks following adoptive transfer in purified CAR T-cells and at one month in whole blood (**table S1**). Lentiviruses favor integration into actively transcribed sites (*15*), and tend to integrate within the gene body rather than near promoters, suggesting low risk for oncogenic transformation (*16*). Indeed, of the integration events we identified within 50kB of *TET2*, ∼94% of CLL sites and ∼93% of ALL sites occurred within the transcriptional boundary of the *TET2* transcriptional unit (**fig. S1D**). The repeated lentiviral integration of a CAR transgene into *TET2* motivates deeper analysis of the role of TET2 in T-cell biology and especially in CAR T-cell differentiation.

### *TET2*-deleted CAR T-cells adopt a central memory-like state post manufacturing

*TET2* loss correlates with clinical response to CAR T-cell therapy (*10*) and its deletion promotes acquisition of a memory CD8^+^ T-cell fate in the setting of acute infection (*14*). To investigate the impact of TET2 loss in human CAR T-cell differentiation, we generated TET2-deficient CAR T-cells (*TET2_KO_*) through CRISPR/Cas9 gene-editing (**Fig. 1A and fig. S2, A-C**). Following lentiviral CAR transduction and primary expansion, we observed a slightly elevated proportion of central memory CAR T-cells in the TET2-deficient condition (**Fig. 1B**). Mitochondrial respiration profiling indicated that, post-production, *TET2_KO_* CAR T-cells were programmed for enhanced oxidative phosphorylation (**Fig. 1C**), with increased basal respiration, maximal respiration, and spare respiratory capacity (SRC) (**Fig. 1D**). *TET2_KO_* CAR T-cells also had increased aerobic glycolysis (**Fig. 1E**). Thus, TET2 deficiency augments cellular metabolism, potentially providing a greater ATP reserve during heightened cellular activity or metabolic stress, aligning with the bioenergetic advantage and rapid recall capability of memory CD8^+^ T-cells (*17*).

**Figure 1.**
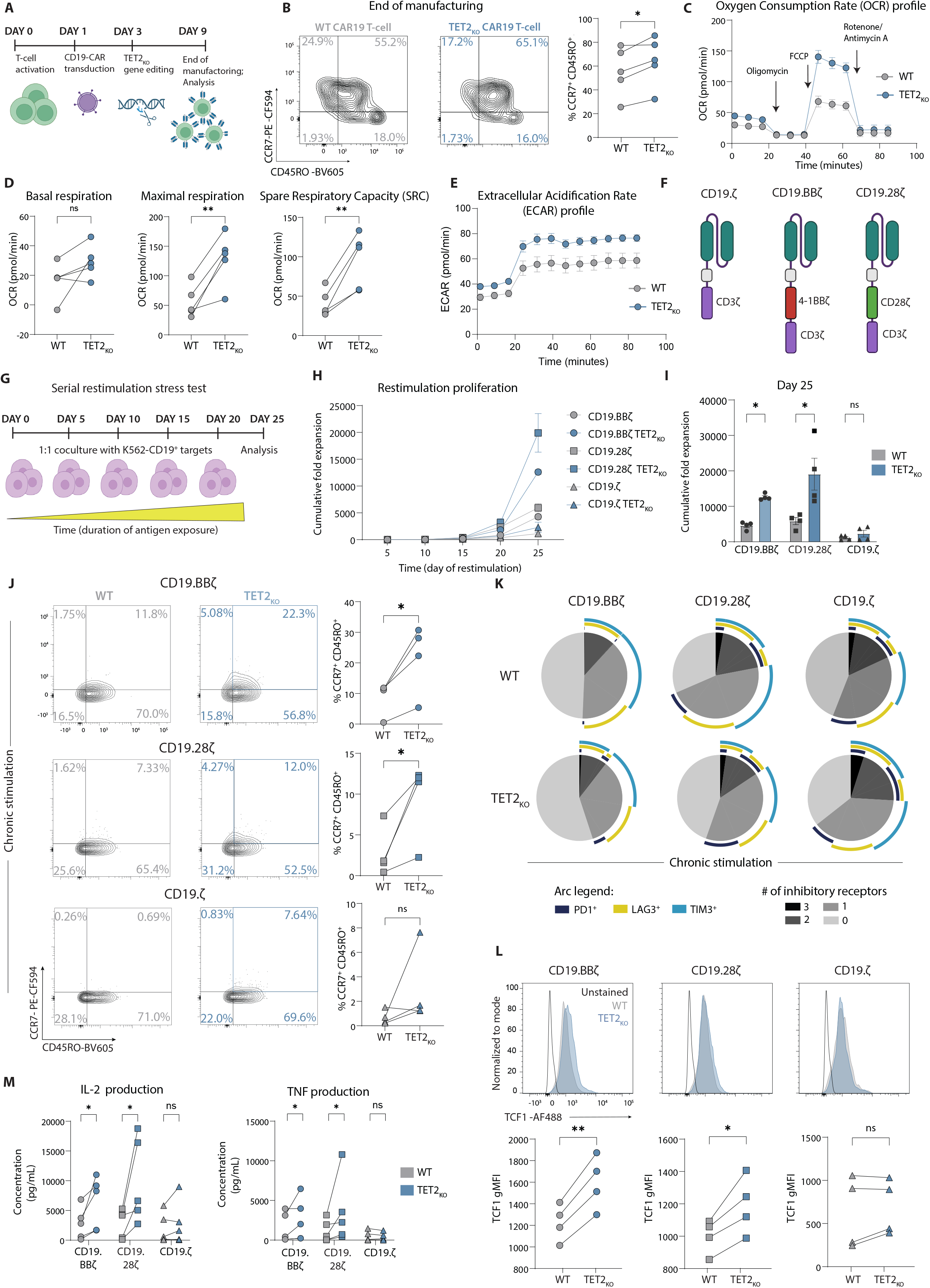
TET2-deleted CAR T-cells adopt a central memory-like state following manufacturing and exhibit increased expansion and reduced IR expression under chronic stimulation across multiple CAR constructs. (**A**) Schematic of CD19 CAR T-cell manufacturing and *TET2* gene editing. (**B**) Example plots and data after CAR T-cell expansion (day 9), highlighting CCR7^+^ CD45RO^+^ central memory subset, n = 5. (C-E) Oxygen Consumption Rate (OCR) (**C**), Basal respiration, maximal respiration, Spare Respiratory Capacity (SRC) (**D**) and Extracellular Acidification Rate (ECAR) (**E**) of *TET2*-disrupted cells at end of expansion (day 9) after administration of oligomycin, FCCP and antimycin A/rotenone as indicated by arrows in (C), n = 5, run in triplicate. (**F-G**) Schematic of CD19.CD3ζ, CD19.BBζ, and CD19.CD28ζ CAR constructs ± *TET2_KO_* (**F**) placed in a serial restimulation stress test (**G**). (**H-I**) Cumulative fold expansion of serially restimulated CAR T-cells ± *TET2_KO_* throughout restimulation (H) and at day 25 (**I**), n = 4. (**J**) Example plots and data of serially restimulated CAR T-cells ± *TET2_KO_* showing distribution of CCR7^+^ CD45RO^+^ central memory-associated markers in CD8^+^ CAR T-cell populations after 5 stimulations, n = 4. (**K**) SPICE plots showing distribution of inhibitory receptor co-expression in serially restimulated CD8^+^ CAR T-cells ± *TET2_KO_* after 5 stimulations, n = 4. (**L**) Example histograms (top) and data (bottom) of TCF1 gMFI in serially restimulated CD8^+^ CAR T-cells ± *TET2_KO_* after 5 stimulations, n = 4. (**M**) IL-2 and TNF production from supernatant collected 24 hours after 5^th^ stimulation, n = 5. Data shown as mean ± SEM (**C, E, H**) or individual values (**B, D, I, J, L, M**) from independent donors. ns p > 0.05; * p < 0.05; ** p < 0.01 by paired t-test.

### TET2 loss increases expansion and reduces IR expression after chronic stimulation for multiple CAR constructs

CD8^+^ T-cell exhaustion is characterized by bioenergetic insufficiencies and altered glycolysis (*18, 19*). The metabolic profile of *TET2_KO_* CAR T-cells suggested that targeting *TET2* might improve CD8^+^ T cell survival and function in the setting of chronic antigen stimulation. To examine the role of TET2 in the long-term persistence of CAR T-cells and to investigate the role of specific co-stimulatory domains (**Fig. 1F**), we used an *in vitro* “stress test” incorporating chronic antigen stimulation that recapitulates several features of progressive T-cell exhaustion (**Fig. 1G**). Both *TET2_KO_* 41BB-costimulated CAR T-cells (CD19.BBζ) and *TET2_KO_* CD28-costimulated CAR T-cells (CD19.28ζ) demonstrated greater proliferative capacity following repeated antigen stimulation when compared to *AAVS1_KO_* control CAR T-cells and CAR T-cells lacking a costimulatory domain (CD19.ζ) (**Fig. 1H and I**). Given this increased proliferative capacity, we next investigated the differentiation and phenotype of chronically stimulated *TET2*_KO_ CAR T-cells. After chronic stimulation, CD8^+^ *TET2_KO_* CAR T-cells skewed towards a CCR7^+^ CD45RO^+^ central memory-like population (**Fig. 1J, fig. S2D**). High IR expression and decreased expression of the transcription factor (TF) TCF1 is associated with terminal differentiation of CD8^+^ T-cells (*20*). However, CD8^+^ *TET2_KO_* CAR T-cells had decreased co-expression of IRs including PD1 and TIM3 (**Fig. 1K**) and increased expression of TCF1 (**Fig. 1L**) compared to *AAVS1_KO_* control CAR T-cells, suggesting that, in the absence of TET2, CAR T-cells were less terminally differentiated. Moreover, TET2 loss resulted in increased IL-2 and TNF production from the total CAR T-cell product following overnight re-stimulation with tumor cells after chronic antigen stimulation (**Fig. 1M**). Finally, TET2 knockout similarly impacted the phenotype of both 41BB- and CD28-costimulated CAR T-cells, suggesting that the role of TET2 is independent of the co-stimulatory domain used. Together, these data implied that loss of TET2 restrained CAR T-cell terminal differentiation during chronic antigen stimulation.

### TET2 loss enhances CAR T-cell efficacy in a tonic CAR signaling model

To further investigate the potential role of TET2 in CAR T-cell terminal differentiation, we next knocked out *TET2* in HA.28ζ-CAR T-cells (**fig. S2E**), which exhibit robust tonic signaling and attain functional, transcriptomic and epigenetic features of exhaustion by day 11 of culture (*3*). Expression of progenitor/stem cell-associated markers CCR7 (**fig. S2F**), CD27 and CD62L (**fig. S2G**) were increased on *TET2_KO_* compared to *AAVS1_KO_* HA.28ζ-CAR T-cells, supporting the idea that *TET2_KO_* CD8^+^ T-cells are less terminally differentiated. Next, we tested whether the phenotypic reprogramming of exhausted CAR T-cells induced by TET2 loss would confer enhanced efficacy. *TET2_KO_* HA.28ζ-CAR T-cells exhibited superior expansion (**fig. S2H**), increased cytotoxicity by the bulk CAR T-cell product (**fig. S2I-J**), and enhanced cytokine secretion (**fig. S2K**) compared to *AAVS1_KO_* HA.28ζ-CAR T-cells when co-cultured with 143B-GL osteosarcoma or with NALM-6-GD2 leukemia (**fig. S2, L-M**) cells. Together, these data indicate that *TET2* deletion may improve CAR T-cell efficacy in the setting of chronic antigen exposure and tonic CAR signaling, potentially by limiting terminal differentiation and increasing expression of proteins involved in T-cell survival/persistence.

### TET2 mediates the transition out of the T_EX_ progenitor pool and towards terminal exhaustion in chronic viral infection

In CAR T-cell exhaustion models using chronic antigen stimulation or tonic antigen receptor signaling, *TET2* disruption limited acquisition of some features of exhaustion, such as IR expression, and enriched for expression of proteins associated with cell renewal, including TCF1. Despite the utility of CAR T-cell exhaustion models, the distinct and complex developmental trajectory of CD8^+^ T-cell exhaustion is likely incompletely recapitulated in reductionist *in vitro* systems (*21*). Furthermore, CAR T-cell systems require T-cell activation for CRISPR-Cas9-mediated *TET2_KO_* and CAR transduction, preventing the study of TET2-deficient T_EX_ generated from naïve T-cells. To further investigate the role of TET2 in the developmental trajectory of T_EX_, we used the well characterized LCMV clone 13 chronic infection model (*22–25*). TET2^fl/fl^ CD4^Cre+^ mice were crossed with T-cell receptor (TCR) transgenic P14 mice that express a TCR specific for LCMV D^b^GP^33-41^ to generate *TET2_KO_* P14 mice (TET2^fl/fl^ CD4^Cre+^ P14). *TET2_KO_* P14 cells were adoptively co-transferred with WT P14 cells at a 1:1 ratio into recipient mice. Recipient mice were then infected with LCMV clone 13 and co-transferred P14 cells were analyzed throughout chronic infection (**Fig. 2A and fig. S3A**).

**Figure 2.**
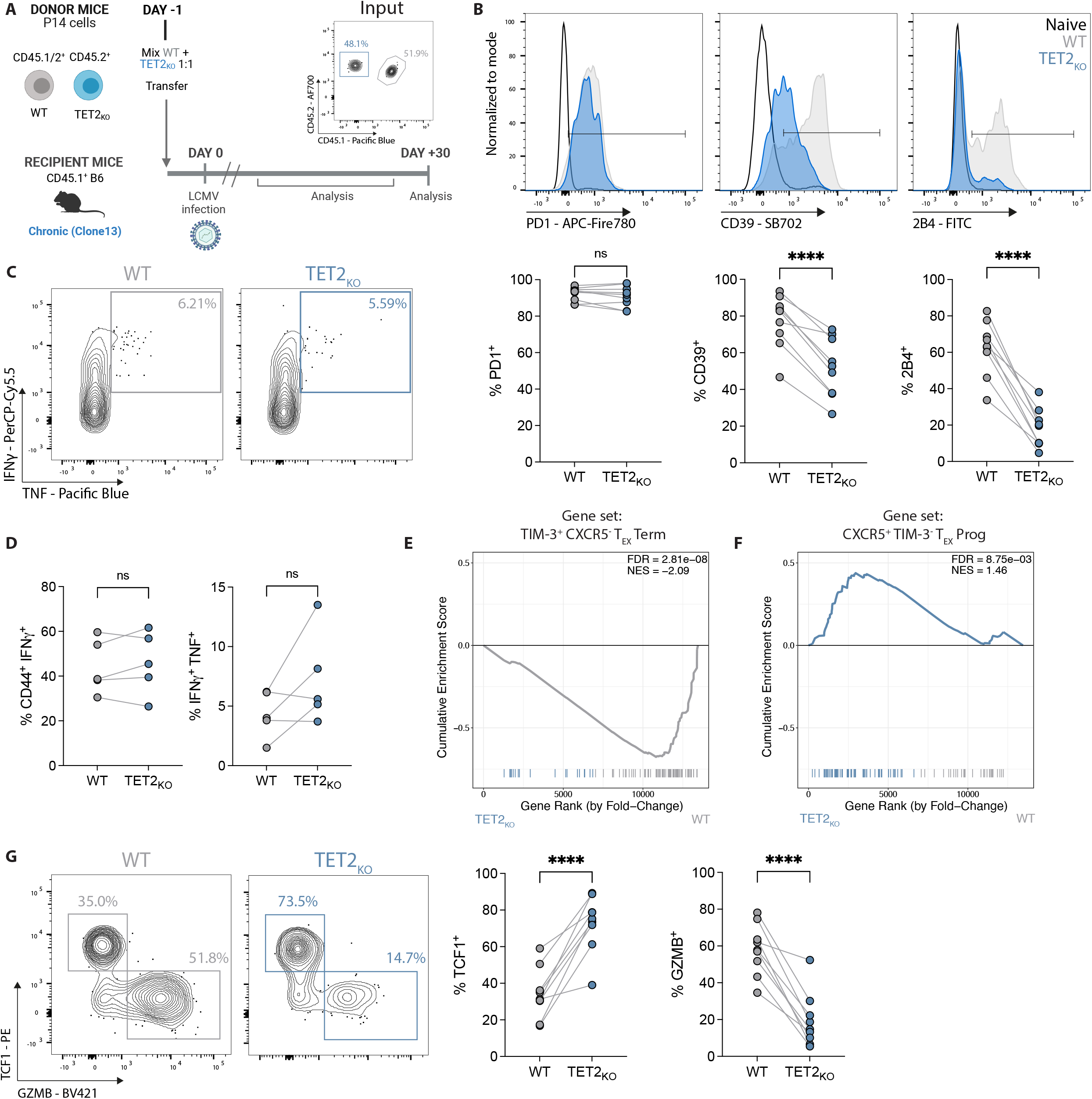
TET2 mediates the transition out of the progenitor T_EX_ subset towards terminal exhaustion.=. (**A**) Co-transfer experimental schematic. Inset plot shows initial P14 co-transfer. (**B**) Example plots and data for inhibitory receptor expression on *TET2_KO_* P14 cells compared to WT P14 cells. (**C-D**) Example plots (**C**) and data (**D**) comparing IFNγ and TNF expression following peptide restimulation for WT and *TET2_KO_* P14 cells. (**E-F**) GSEA of terminal T_EX_ (**E**) and T_EX_ progenitor (**F**) signatures between WT and *TET2_KO_* P14 cells at day 15 p.i. with LCMV clone 13 (Gene sets from GSE84105). (**G**) Example plots and data comparing TCF1^+^ T_EX_ progenitor and GZMB^+^ terminally differentiated T_EX_ frequencies within WT and *TET2_KO_* P14 cells. (**B, G**) n = 9, spleen at day 30 p.i. with LCMV clone 13. Data for individual mice shown; representative of >4 independent experiments. (**D**) n = 5, spleen at day 37 p.i. with LCMV clone 13. Data for individual mice shown; representative of 3 independent experiments. (**B, D, G**) ns p > 0.05; * p < 0.05; ** p < 0.01; *** p < 0.001; **** p < 0.0001 by paired t-test.

The impact of TET2 loss on frequencies of antigen specific CD8^+^ T-cells over the course of chronic infection was variable and influenced by factors in the LCMV model. For example, when recipient wild-type CD4^+^ T-cells were present (i.e., no CD4^+^ T-cell depletion prior to infection), WT P14 outnumbered *TET2_KO_* P14 cells by ∼7:1 in blood during the early stage of chronic infection (∼day 8 post-infection, p.i., **fig. S3B**), despite initial transfer at a 1:1 ratio. However, once exhaustion was established (*26*) *TET2_KO_* P14 cells expanded, outcompeting WT P14 cells by ∼1.8:1 at ∼day 60 pi (**fig. S3B**) and reflecting the expansion seen in human CAR T-cell models. In contrast, when CD4^+^ T-cells were depleted by *in vivo* administration of anti-CD4 antibody GK1.5, *TET2_KO_* P14 cells often did not rebound and remained underrepresented compared to WT P14 cells in blood throughout infection (**fig. S3C**). Furthermore, despite underrepresentation in blood, the frequency of *TET2_KO_* P14 cells in spleen was often more comparable to that of WT P14 cells (**fig. S3D**). Together, these data suggest that the role of TET2 in T_EX_ proliferation/survival may be impacted by CD8^+^ T-cell extrinsic pressures that regulate exhaustion, such as chronic antigen burden/vial load and CD4^+^ T-cell help.

CD8^+^ T-cell exhaustion is characterized by high IR expression and decreased production of effector cytokines (reviewed in (*1*)). We first asked whether *TET2_KO_* P14 cells retained core features of exhaustion in the LCMV chronic infection model. Expression of some IRs such as PD1 (**Fig. 2B**) and LAG3 (**fig. S3E**) remained high on *TET2_KO_* P14 cells and was comparable to co-transferred WT P14 cells. However, TET2 loss strongly decreased expression of other IRs including CD39 and 2B4 (**Fig. 2B**). For example, at day 30 p.i. in spleen, only ∼20% of *TET2_KO_* P14 cells expressed 2B4 compared to ∼63% of WT P14 (**Fig. 2B**). However, *TET2_KO_* P14 cells did not acquire surface characteristics of classical T_EFF_ and T_MEM_ that arise during acute resolving infections (*27, 28*). Rather, *TET2_KO_* KLRG1^+^ T_EFF_-like P14 cells were effectively absent from the spleen at day 30 p.i., whereas WT KLRG1^+^ P14 cells were detectable at low frequencies as expected (**fig. S3F-G**). Although *TET2_KO_* P14 cells had moderately increased expression of the IL-7 receptor CD127, which is associated with memory-like differentiation, protein levels remained low and consistent with expression in chronic rather than acute infection (**fig. S3F-G**). Furthermore, despite decreased expression of some key IRs, *TET2_KO_* P14 cells remained functionally exhausted, as a similar frequency of *TET2_KO_* P14 cells produced IFNγ or co-produced IFNγ and TNF following *in vitro* restimulation with LCMV peptide as WT P14 cells (**Fig. 2, C and D**).

To investigate transcriptional changes associated with loss of TET2 during chronic infection, we next performed bulk RNA-sequencing on WT and *TET2_KO_* P14 cells at day 15 p.i. with LCMV clone 13. *TET2_KO_* P14 cells had a distinct transcriptional profile, with ∼1750 genes differentially expressed between WT and *TET2_KO_* P14 cells (FDR<0.05, **fig. S3H, table S3**), including decreased expression of the IRs *Entpd1* (CD39) and *Cd244* (2B4), supporting protein expression data. The T_EX_ lineage is functionally diverse. T_EX_ progenitor cells retain proliferative potential, express TCF1, and have decreased expression of specific IRs such as CD39 and TIM3 (*29–35*) despite high expression of the exhaustion-associated TF TOX (*36–41*). T_EX_ progenitors differentiate into a terminal T_EFF_-like subset that have reacquired some effector functions and contribute to viral control (*35, 42–46*) or into terminally exhausted T_EX_ cells with increased IR expression and decreased proliferative capacity (*43*). Loss of TET2 decreased expression of genes associated with terminal differentiation and effector biology, including *Zeb2* (*47, 48*), *Id2* (*49*), *Klrg1* (*50, 51*), *Nkg7* (*52, 53*), *Gzma* and *Gzmk* (*54*) (**fig. S3H**). The loss of effector TFs and molecules, along with decreased expression of certain IRs, as well as our initial CART-cell data, provoked the hypothesis that TET2 may have a role in terminal T_EX_ differentiation. Gene Set Enrichment Analysis (GSEA) revealed depletion of a terminally differentiated T_EX_ gene set in *TET2_KO_* P14 cells (**Fig. 2E**), suggesting a decrease in terminal exhaustion compared to WT controls. Conversely, a T_EX_ progenitor gene set was enriched in *TET2_KO_* P14 cells (**Fig. 2F**); indeed, genes associated with T_EX_ progenitor biology including *Slamf6* (LY108) and *Tnfsf4* (OX40L) were upregulated when *TET2* was knocked out (**fig. S3H**). These gene expression differences were associated with robust changes in T_EX_ subset distribution. *TET2_KO_* P14 cells had a relative increase in the proportion of T_EX_ progenitor cells (TCF1^+^ GZMB^−^) with a marked reduction in terminally differentiated T_EX_ (TCF1^−^ GZMB^+^) (**Fig. 2G**). These observations suggest that TET2 regulates differentiation into terminally differentiated T_EX_ subsets, including effector-like T_EX_ cells, during chronic infection.

### Loss of TET2 limits terminal differentiation of exhausted CD8^+^ T-cells

Bulk RNA-seq comparing WT and *TET2_KO_* P14 cells suggested that TET2 acts at a checkpoint in the epigenetic transition between T_EX_ progenitor and terminally differentiated T_EX_ cell fates, driving terminal exhaustion at the expense of a stem cell-like state. To determine if TET2 functions at the transition between T_EX_ subsets, or regulates T_EX_ differentiation within these subsets, we isolated LY108^+^ T_EX_ progenitor and LY108^neg^ terminally differentiated T_EX_ from WT and *TET2_KO_* P14 cells at day 15 p.i. with LCMV clone 13 and performed RNA-seq and Assay for Transposase Accessible Chromatin Sequencing (ATAC-seq) on the isolated subsets (**Fig. 3, A and B**). T_EX_ progenitor and terminally differentiated T_EX_ cells are transcriptionally and epigenetically distinct (*34*). Principal component analysis (PCA) revealed that cell subset (LY108^+^ versus LY108^neg^ T_EX_) was the major contributor to sample-to-sample variation and separated samples along PC1 regardless of genotype (**Fig. 3, C and D**). In contrast, PC2 was driven by genotype, with all isolated *TET2_KO_* P14 populations localized in distinct regions compared to WT P14 subsets (**Fig. 3, C and D**). Therefore, although *TET2_KO_* T_EX_ cells retain key features of WT T_EX_ (**Fig 2. B-D**, **fig. S3H**), loss of TET2 may impact differentiation within T_EX_ subsets.

**Figure 3.**
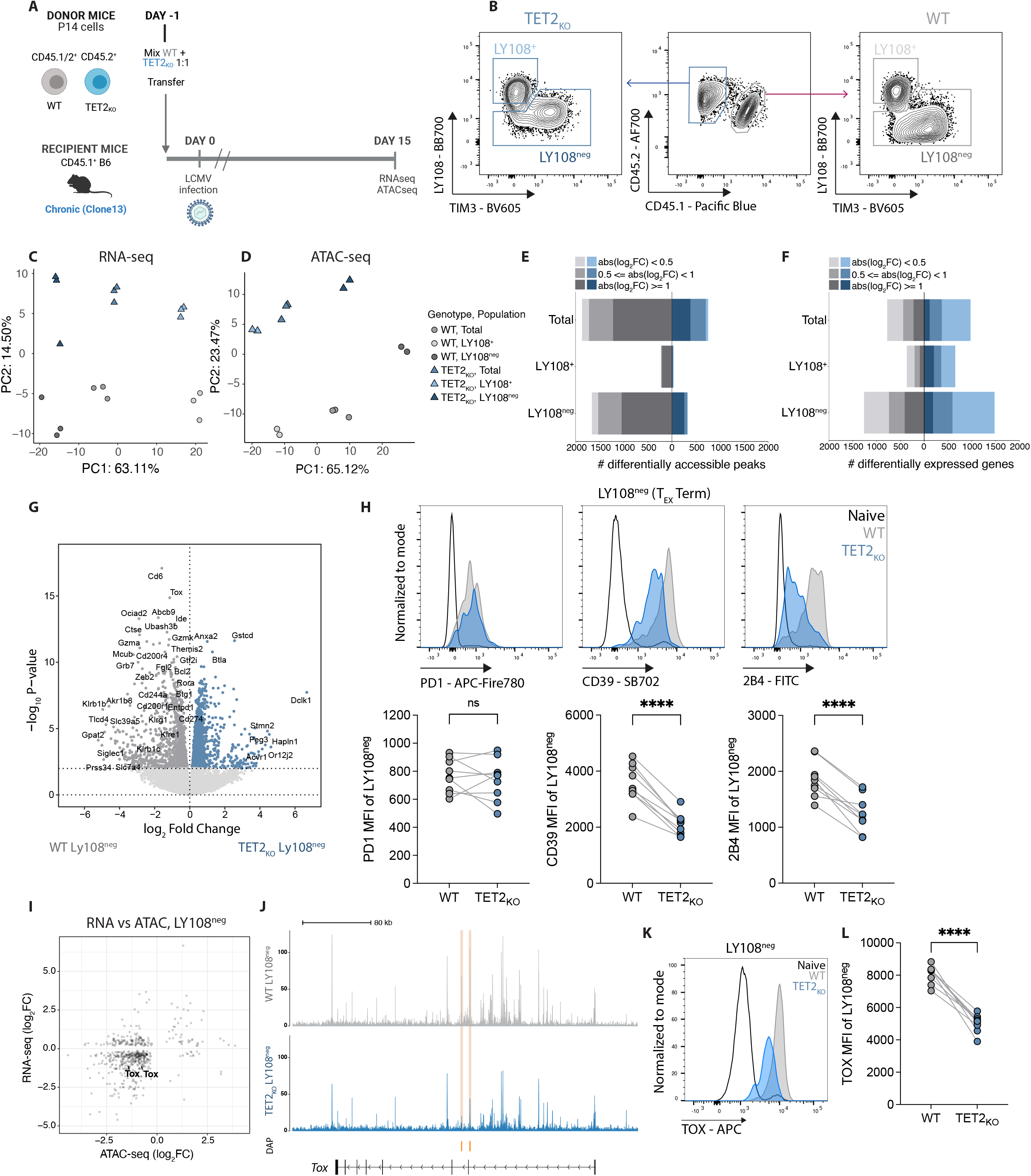
Loss of TET2 limits terminal differentiation of exhausted CD8 T-cells. (**A**) Experiment schematic for RNA and ATAC sequencing. WT and *TET2_KO_* P14 cells were analyzed at day 15 p.i. with LCMV clone 13. (**B**) Sorting strategy for T_EX_ subsets for RNA and ATAC sequencing. (**C-D**) PCA of RNA sequencing (**C**) and ATAC sequencing (**D**) data for WT and *TET2_KO_* T_EX_ subsets. (**E-F**) Number of DACRs (**E**) or DEGs (**F**) for each pairwise comparison between WT and *TET2_KO_* T_EX_ subsets (FDR < 0.05, with variable absolute log_2_ fold changes (abs(log_2_FC)) indicated). (**G**) Volcano plot highlighting DEG in WT compared to *TET2_KO_* LY108^neg^ T_EX_ cells. (**H**) Example plots and data comparing expression of PD1, CD39 and 2B4 on *TET2_KO_* LY108^neg^ T_EX_ to WT LY108^neg^ T_EX_. (**I**) Correlation plot of differential gene expression and peak accessibility in *TET2_KO_* LY108^neg^ T_EX_ compared to WT LY108^neg^ T_EX_ with TOX labelled. (**J**) Example tracks showing accessibility at the *Tox* locus in LY108^neg^ T_EX_. Differentially Accessible Peaks (DAPs) are indicated in orange. (**K-L**) Example plots (**K**) and data (**L**) comparing TOX expression in *TET2_KO_* LY108^neg^ T_EX_ to WT LY108^neg^ T_EX_. (**H, L**) n = 9, spleen at day 30 p.i. with LCMV clone 13. Data for individual mice shown; representative of 3 independent experiments. ns p > 0.05; **** p < 0.0001 by paired t-test.

To examine the potential role of TET2 within T_EX_ subsets, we directly compared WT and *TET2_KO_* T_EX_ within isolated T_EX_ subsets and identified differentially expressed genes (DEG) and differentially accessible chromatin regions (DACR). These analyses revealed two major features of TET2 function in T_EX_ differentiation. First, 1855 DACR and 2744 DEG distinguished WT from *TET2_KO_* LY108^neg^ terminally differentiated T_EX_ compared to only 255 DACR and 1018 DEG between WT and *TET2_KO_* LY108^+^ T_EX_ progenitors (**Fig. 3, E and F**; **fig. S4, A and B, table S3 and S4**). Reflecting these relative differences, WT and *TET2_KO_* LY108^+^ T_EX_ progenitors were closer to each other in PCA space than WT and *TET2_KO_* terminally differentiated T_EX_ (**fig. S4, C and D**), indicating that terminally differentiated T_EX_ cells are more transcriptionally and epigenetically distinct following loss of TET2 than the T_EX_ progenitor subset. Second, within the terminally differentiated T_EX_ subset, the majority of DACR (82.7%; FDR<0.05, log2 fc > 0.5) were less accessible following *TET2* knockout. Together, these data suggest that TET2 is required to sustain and/or increase chromatin accessibility at, and expression of, genes associated with terminal T_EX_ differentiation.

We next asked which genes were unable to be upregulated in LY108^neg^ terminally differentiated T_EX_ in the absence of TET2. Multiple genes associated with effector functions, including KLR family members (*Klrg1, Klrb1b, Klrb1c and Klre1)*, cytotoxic markers *Gzma* and *Gzmk* and the TF-encoding genes *Zeb2, Btg1* and *Rora* were decreased in expression in *TET2_KO_* compared to WT terminally differentiated T_EX_ (**Fig. 3G**). Furthermore, TF motif analysis revealed that binding sites for T_EFF_-associated TFs including RUNX and TBET were less accessible in terminally differentiated T_EX_ following removal of TET2 (**fig. S4E**). Thus, in terminally differentiated T_EX_ cells lacking TET2, binding sites for key effector-driving TFs are less accessible and expression of effector genes is diminished. These data support the notion that TET2 promotes the reacquisition of effector-associated genes in the terminally differentiated T_EX_ subset.

RNA and protein expression of key IRs including CD39 (*Entpd1*) was lower on total *TET2_KO_* P14 cells than WT P14 cells (**Fig. 2B**, **S3H**). To determine if this change in IR expression reflected the population shift towards T_EX_ populations with decreased IR expression (T_EX_ progenitor cells) or differential regulation of IRs within terminally differentiated subsets, we next assessed IR expression within isolated T_EX_ subsets. Inhibitory receptor expression, including *Cd200r1* (CD200R), *Entpd1* (CD39), *Cd274* (PDL1) and *Cd244* (2B4), was decreased in *TET2_KO_* terminally differentiated T_EX_ (**Fig. 3G and H**), whereas levels of these IRs were low and more comparable to WT controls for *TET2_KO_* T_EX_ progenitors (**fig. S4F and G**). LY108 (*Slamf6*) decreases in expression as T_EX_ terminally differentiate (*34*); however, LY108 expression remained high in *TET2_KO_* terminally exhausted T_EX_ cells (**fig. S4H**). Together, these data support the hypothesis that TET2 regulates loss of T_EX_ progenitor biology and is required for complete differentiation within the terminally exhausted T_EX_ population.

The transcription factor TOX has been proposed to regulate terminal exhaustion (*34*). Therefore, we interrogated if the decreased differentiation with the terminal T_EX_ subset following *TET2_KO_* was associated with changes in TOX. Indeed, *Tox* expression was decreased within *TET2_KO_* LY108^neg^ terminal T_EX_ and this decreased expression was associated with reduced chromatin accessibility at the *Tox* locus (**Fig. 3, I and J**). These changes in RNA expression and chromatin accessibility translated to markedly reduced TOX protein expression in *TET2_KO_* terminal T_EX_ compared to WT terminal T_EX_ (**Fig. 3, K and L**). Therefore, TET2 may coordinate with TOX to regulate the terminal differentiation of T_EX_.

### TET2 regulates early bifurcation of T_EX_ from T_EFF_-like cells

At least three major epigenetic remodeling events underpin the developmental trajectory of T_EX_ cells. The first occurs immediately following naïve CD8^+^ T-cell activation. The second occurs early, within days of initial activation, when terminally differentiated T_EFF_-like cells bifurcate from TCF1^+^ T_EX_ precursors. Analysis of *TET2_KO_* P14 cells late in chronic infection suggested that TET2 regulates the third major rewiring event occurring in established T_EX_, when T_EX_ progenitors transition into effector-like and terminally exhausted T_EX_ subsets (*34*). RNA-seq and ATAC-seq analysis of terminally differentiated T_EX_ cells suggested a role in coordinating the re-acquisition of effector-like biology. To interrogate the role of TET2 in the second bifurcation event prior to fate-commitment to exhaustion, we set up an adoptive co-transfer of WT and *TET2_KO_* P14 cells as described above, then analyzed CD8^+^ T-cell responses to chronic LCMV infection at early timepoints (**Fig. 4A**). At day 6 and 8 p.i. the proportion and absolute frequency of PD1^low^ KLRG1^+^ T_EFF_ were markedly reduced in the absence of TET2 (**Fig. 4, B and C**, **fig. S4I**). Furthermore, *TET2_KO_* P14 cells skewed towards TCF1^+^ T_EX_ precursors at the expense of granzyme B-expressing T_EFF_-like cells (**Fig. 4D**). Together, these findings imply that TET2 regulates the acquisition of effector-like biology at multiple steps in T_EX_ differentiation, both in the early bifurcation between T_EX_ precursors and T_EFF_ and in established exhaustion.

**Figure 4.**
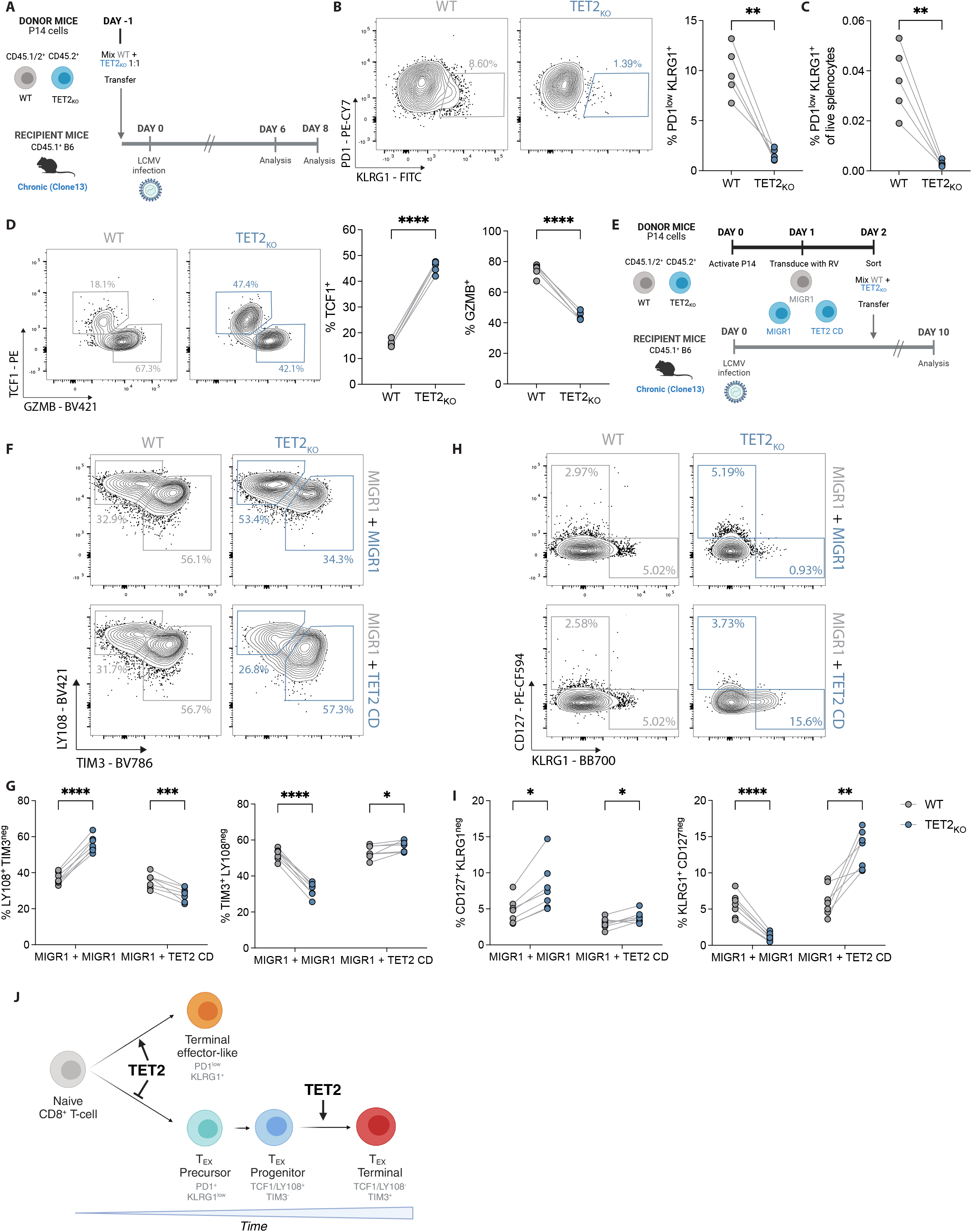
TET2 regulates early bifurcation of T_EX_ from T_EFF_-like cells. (**A**) Experiment schematic for analysis of *TET2_KO_* P14 cells early (day 6-8) of LCMV clone 13 infection. (**B**) Example plots and data comparing frequency of early T_EFF_-like cells (KLRG1^+^ PD1^low^) within WT and *TET2_KO_* P14 cells at day 8 p.i. (**C**) Frequencies of WT and *TET2_KO_* KLRG1^+^ PD1^low^ T_EFF_-like cells from total live splenocytes at day 8 p.i. (**D**) Example plots and data comparing expression of TCF1 and GZMB for WT and *TET2_KO_* P14 cells. (**E**) Experiment schematic for rescue of TET2 function in *TET2_KO_* P14 cells. (**F-G**) Example plots (**F**) and data (**G**) comparing frequencies of LY108^+^ T_EX_ and TIM3^+^ T_EX_ for WT and *TET2_KO_* P14 cells with or without overexpression of the TET2 catalytic domain (TET2 CD versus MIGR1). (**H-I**) Example plots (**H**) and data (**I**) comparing frequencies of CD127^+^ T_MEM_-like and KLRG1^+^ T_EFF_-like cells for WT and *TET2_KO_* P14 cells with or without overexpression of the TET2 catalytic domain (TET2 CD). (**J**) Model for TET2 role at major bifurcation events in chronic infection. (**B, C, D**) n = 5, spleen at day 8 p.i. with LCMV clone 13. Data for individual mice shown, representative of 2 independent experiments. * p < 0.05; ** p < 0.01; *** p < 0.001; **** p < 0.0001 by paired t-test. (**G, I**) n = 7, spleen at day 10 p.i. with LCMV clone 13. Data for individual mice shown, representative of 2 independent experiments. ns p > 0.05; * p < 0.05; ** p < 0.01; *** p < 0.001; **** p < 0.0001 by multiple paired t-test with Holm-Šídák post-test correction.

These data suggested that TET2-deficiency limits acquisition of effector T-cell-like biology. To test whether forced TET2 expression rescues the phenotypes observed *TET2_KO_* P14 cells or promotes T-cell effector biology, we overexpressed the TET2 catalytic domain (TET2 CD) in *TET2_KO_* P14 and compared the impact of TET2 “rescue” to *TET2_KO_* P14 cells, as well as WT P14 cells transduced with an empty vector (MIGR1) (**Fig. 4E**). As previously observed, T_EX_ subset distribution in *TET2_KO_* P14 was skewed towards T_EX_ progenitors at the expense of terminal T_EX_ differentiation (**Fig. 4, F and G**). In contrast, expression of the TET2 catalytic domain normalized the proportions of T_EX_ subsets and pushed T_EX_ slightly towards terminal differentiation (**Fig. 4, F and G**). Furthermore, whereas KLRG1^+^ *TET2_KO_* P14 cells were effectively absent, expression of the TET2 catalytic domain increased KLRG1 expression by ∼2-fold compared to WT P14 cells and ∼11.6-fold compared to *TET2_KO_* P14 cells (**Fig. 4, H and I**). Therefore, TET2 promotes terminal T_EX_ differentiation and the TET2 catalytic domain is sufficient for this activity.

Together, these data imply that TET2 acts a rheostat to regulate terminal differentiation and the acquisition of effector-like biology at multiple checkpoints in CD8^+^ T-cell development (**Fig. 4J**). In acute infection, TET2 modulates the bifurcation between classical short-lived T_EFF_ and memory precursors (*14*), enforcing terminal differentiation and the acquisition of effector functions. TET2 plays parallel roles in chronic infection. In the initial stages of chronic infection, TET2 drives CD8^+^ T-cells towards T_EFF_-like cells and away from the formation of the T_EX_ precursor pool (*55*). Once exhaustion is established, TET2 pushes T_EX_ progenitors towards a terminally differentiated and T_EFF_-like T_EX_ state, again mirroring the role of TET2 as an enforcer of differentiation. These data further provoke the hypothesis that similar epigenetic programs are used and reused throughout CD8^+^ T-cell differentiation to regulate function in distinct contexts (*56*).

### *TET2*-edited dual knock-in allogeneic CAR T-cells resist terminal T_EX_ differentiation, allowing enhanced tumor control

Data from the chronic infection model suggested that TET2 regulates CD8^+^ T-cell differentiation, and that loss of TET2 limits terminal exhaustion. Our initial *TET2*_KO_ CAR T-cell data implied that targeting TET2 to restrain terminal differentiation could improve CAR T-cell expansion and efficacy. However, bi-allelic loss of *TET2* with BATF3 expression led to clonal proliferation of CAR T-cells (*13*) and highlighted the additional considerations required to safely manipulate epigenetic regulators in the clinic. Therefore, we next applied synthetic biology principles to design a clinically actionable CAR T-cell with disrupted TET2 for improved efficacy. A key element of our strategy involved a dual knock-in (KI), simultaneously editing the *TRAC* and *TET2* loci using CRISPR/Cas9 and introducing new genetic templates at these sites. First, we designed a single-guide (sg) RNA to target the 5’ end of the first exon of *TRAC*. This enabled integration of an anti-CD19 CAR from an adeno-associated virus (AAV) donor DNA cassette into the *TRAC* locus and simultaneously resulted in TCR knock-out (**Fig. 5A**). The KI construct (TRAC-CAR19) featured a 41BB co-stimulatory endodomain and co-expressed a truncated nerve growth factor receptor (tNGFR) for selection (**Fig. 5B**). This approach was designed to delay effector T-cell differentiation and exhaustion through CAR insertion at the *TRAC* locus as previously described (*57*), while leveraging the potential benefits of 41BB co-stimulation (*4*). Furthermore, this TCR knockout strategy could enhance therapeutic safety by reducing risks of TCR-induced autoimmunity and alloreactivity. In addition, expression of the TRAC-CAR19 construct is controlled by the endogenous *TRAC* promoter, thus driving physiological CAR expression on the cell surface. The TRAC-CAR19 KI efficiency was proportional to the AAV dosage, achieving over 70% efficiency at a multiplicity of infection (MOI) of 50,000 (**Fig. 5B and fig. S5A**) and between 72-98% of CAR^+^ T-cells were CD3-negative (**Fig. 5C and fig. S5A**), validating this dual knock-out and knock-in strategy.

**Figure 5.**
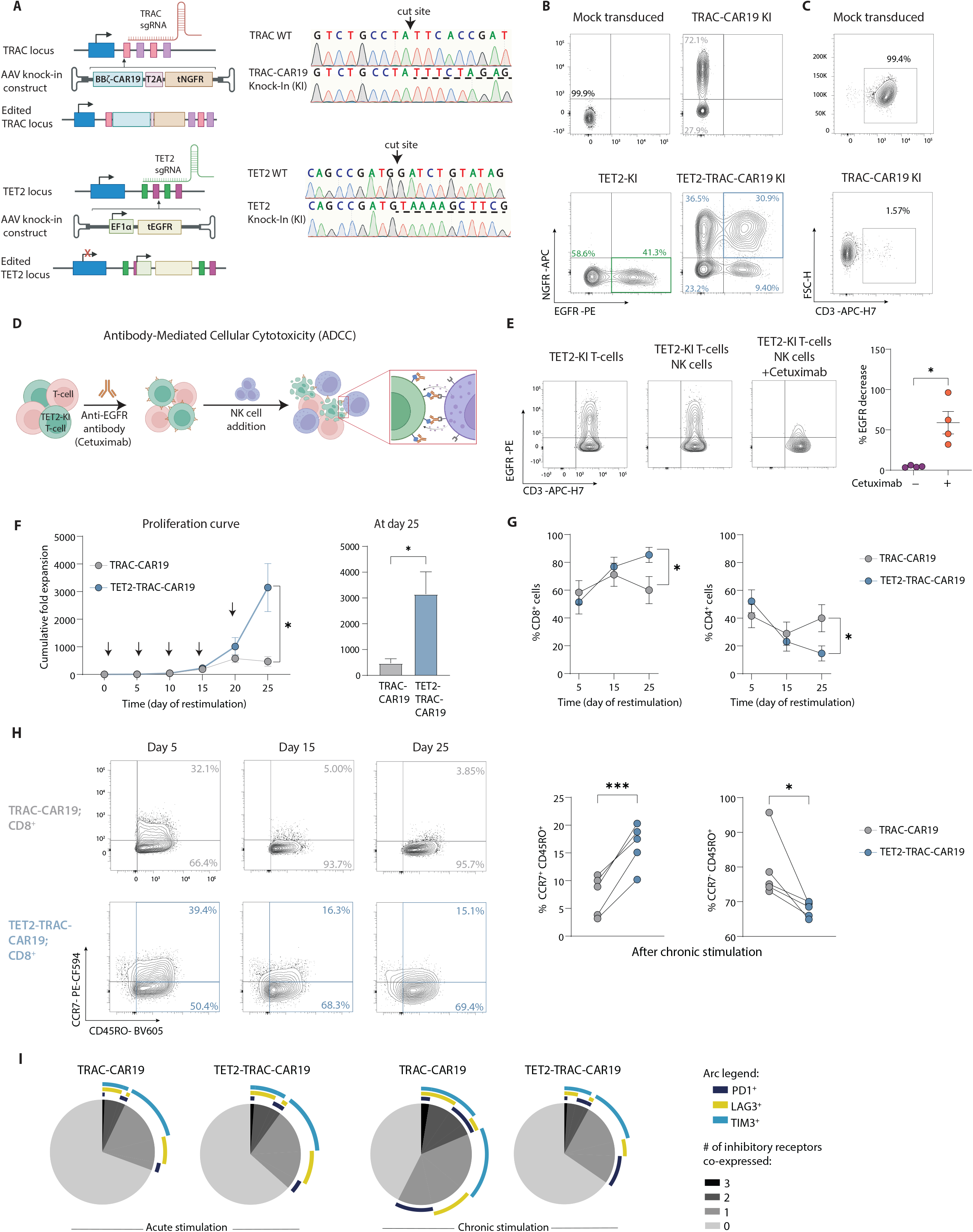
Dual knock-in TET2-TRAC-CAR19 T-cells have enhanced proliferation and maintain memory-associated marker expression. (**A**) Left; Schematic of *TRAC* (**top**) and *TET2* (**bottom**) loci alongside rAAV6 knock-in (KI) vectors. Right; Sanger sequencing electropherogram confirming integration of TRAC and TET2 KI constructs, underlined with dashed line. (**B**) Example plots of TET2 and TRAC-CAR19 single KI or dual TET2-TRAC-CAR19 KI T-cells. (**C**) Example plots of CD3 loss detected by flow in TRAC-CAR19-KI T-cells. (**D-E**) Schematic of *in vitro* ADCC assay (**D**) to deplete CRISPR-edited TET2-KI T-cells. Example plots and data (**E**) of EGFR expression on TET2-KI T-cells alone or in an NK cell co-culture ± Cetuximab incubation, gated on CD56^−^ populations, n = 4. (**F**) Cumulative fold expansion of TRAC-CAR19 and TET2-TRAC-CAR19 T-cells during restimulation and at day 25, arrows represent addition of irradiated K562-CD19^+^ target cells, n = 5. (**G**) Proportions of CD4^+^ versus CD8^+^ T-cells in TRAC-CAR19 and TET2-TRAC-CAR19 T-cells after stimulation, n = 7. (**H**) Example plots showing distribution of central (CCR7^+^ CD45RO^+^) and effector (CCR7^−^ CD45RO^+^) memory-associated markers in CD8^+^ CAR T-cell populations after restimulation, with summary after 5 stimulations, n = 5. (**I**) SPICE plot showing distribution of inhibitory receptor co-expression in CD8^+^ TRAC-CAR19 and TET2-TRAC-CAR19 T-cells after 1 (acute) and 5 (chronic) stimulations, n = 6. (**J**) Data shown as mean ± SEM (**F, G**) or individual values (**E, H**) from independent donors. ns p > 0.05; * p < 0.05; ** p < 0.01; *** p < 0.001 by paired t-test.

In parallel with targeted modifications at the *TRAC* locus, we used CRISPR/Cas9 and a second AAV vector repair matrix to both disrupt *TET2* (TET2-TRAC-CAR19) and integrate a truncated human epidermal growth factor receptor (tEGFR) cDNA at the *TET2* locus (**Fig. 5A and fig. S5A**), under the regulation of an exogenous human EF1α promoter. The successful incorporation and functionality of the tEGFR enabled *in vitro* selection of *TET2*-edited cells **(fig. S5B**). We next tested if tEGFR expression could function as a ‘safety switch’ and allow targeted elimination of *TET2*-disrupted CAR T-cells. TET2-TRAC-CAR19 T-cells were cultured *in vitro* with natural killer (NK) cells and the FDA-approved antibody Cetuximab. Cetuximab targets EGFR and induces antibody-dependent cellular cytotoxicity (ADCC) (**Fig. 5D**). Following co-culture, EGFR-expressing CAR T-cells were selectively depleted (**Fig. 5E**). Thus, tEGFR provides a critical safety switch that allows for controlled depletion of CRISPR-edited cells.

We next confirmed that our dual CRISPR-editing and AAV knock-in CAR T-cell engineering approach did not negatively impact manufacturing. Indeed, TRAC-CAR19 T-cells expanded as expected (**fig. S5C**) and TET2-TRAC-CAR19 T-cells exhibited similar metabolic potency enhancements as *TET2_KO_* CAR T-cells (**fig. S5D**) during manufacturing.

To test if targeting TET2 in TRAC-CAR19 T-cells could provide an advantage in settings of chronic antigen, we isolated edited (tEGFR^+^ and tNGFR^+^; (**fig. S5B**)) TRAC-CAR19 and TET2-TRAC-CAR19 T-cells and subjected them to the *in vitro* restimulation assay described above to recapitulate features of progressive T-cell exhaustion (**Fig. 1G**). During restimulation, TET2-TRAC-CAR19 T-cells demonstrated significant proliferative potential, with a sixfold increase in cumulative expansion by day 25 compared to TRAC-CAR19 T-cells (**Fig. 5F**) that was antigen dependent (**fig. S5E**). This increased expansion only became apparent by the fourth round of stimulation (day 20), suggesting that this proliferative advantage was associated with chronic antigen exposure. The proliferative advantage of *TET2* disruption was most apparent for CD8^+^ T-cells, as CD8^+^ TET2-TRAC-CAR19 T-cells expanded more than the CD4^+^ T-cell equivalent (**Fig. 5G**). Concurrently, TET2-TRAC-CAR19 CD8^+^ T-cells maintained a higher expression of the progenitor-associated receptor CCR7 (**Fig. 5H**) and exhibited lower frequencies of IR co-expression (PD1, LAG3, TIM3) (**Fig. 5I**) than control TRAC-CAR19 CD8^+^ T-cells following chronic antigen stimulation. Together, these data demonstrate that TET2-deficiency improves maintenance of a less terminally differentiated TRAC-CAR19 CD8^+^ T-cell pool under conditions of chronic antigen stimulation, supporting our findings with *TET2_KO_* CAR T-cells and *TET2_KO_* in chronic infection.

To gain a more detailed understanding of the impact of TET2 loss on CAR T-cell responses to chronic antigen stimulation we performed bulk RNA-sequencing on isolated TET2-TRAC-CAR19 CD8^+^ T-cells following four rounds of *in vitro* restimulation. TET2-TRAC-CAR19 cells upregulated progenitor and memory-associated genes typically lowly expressed in terminally exhausted CD8^+^ T-cells including *TCF7*, *CCR7*, *SELL*, and *TOX2* (*58*) (**Fig. 6, A and B, table S5**) compared to control TRAC-CAR19 cells. This increase in expression of progenitor-associated genes was accompanied by a downregulation of genes related to calcium signaling and TF activity including *BRS3, GJA1, CAP2, NANOGNB* and *GBX1* (**Fig. 6A**). GSEA further supported the notion that TET2-TRAC-CAR19 cells retained features of more stem-like cells, with an enrichment for both a T_EX_ progenitor signature and a stem cell/central memory CD8^+^ T cell signature (T_SCM_/T_CM_) **(Fig. 6C**). In contrast, a terminally exhausted tumor-infiltrating lymphocyte signature (*59*) was negatively enriched in TET2-TRAC-CAR19 (**Fig. 6D**). In line with GSEA results, chronic stimulation led to an increase in the proportion of CD8^+^ TET2-TRAC-CAR19 cells expressing the progenitor-associated TF TCF1 (*60*) and a decrease in the proportion of cells expressing the terminal differentiation-associated effector molecule granzyme B (*61*) compared to TRAC-CAR19 control cells (**Fig. 6E, fig. S5F**). Furthermore, TET2-TRAC-CAR19 cells retained the CD8^+^ TCF1^+^ CD62L^+^ population suggested to be essential for T_EX_ progenitor proliferative responses (*62*) (**Fig. 6F, fig. S5G**). Together, these data further suggest that the role of TET2 in chronically stimulated TRAC-CAR19 CD8^+^ T-cells mirrors that identified in chronic LCMV infection, whereby TET2 drives terminal differentiation, and loss of TET2 supports maintenance of a progenitor-like phenotype.

**Figure 6.**
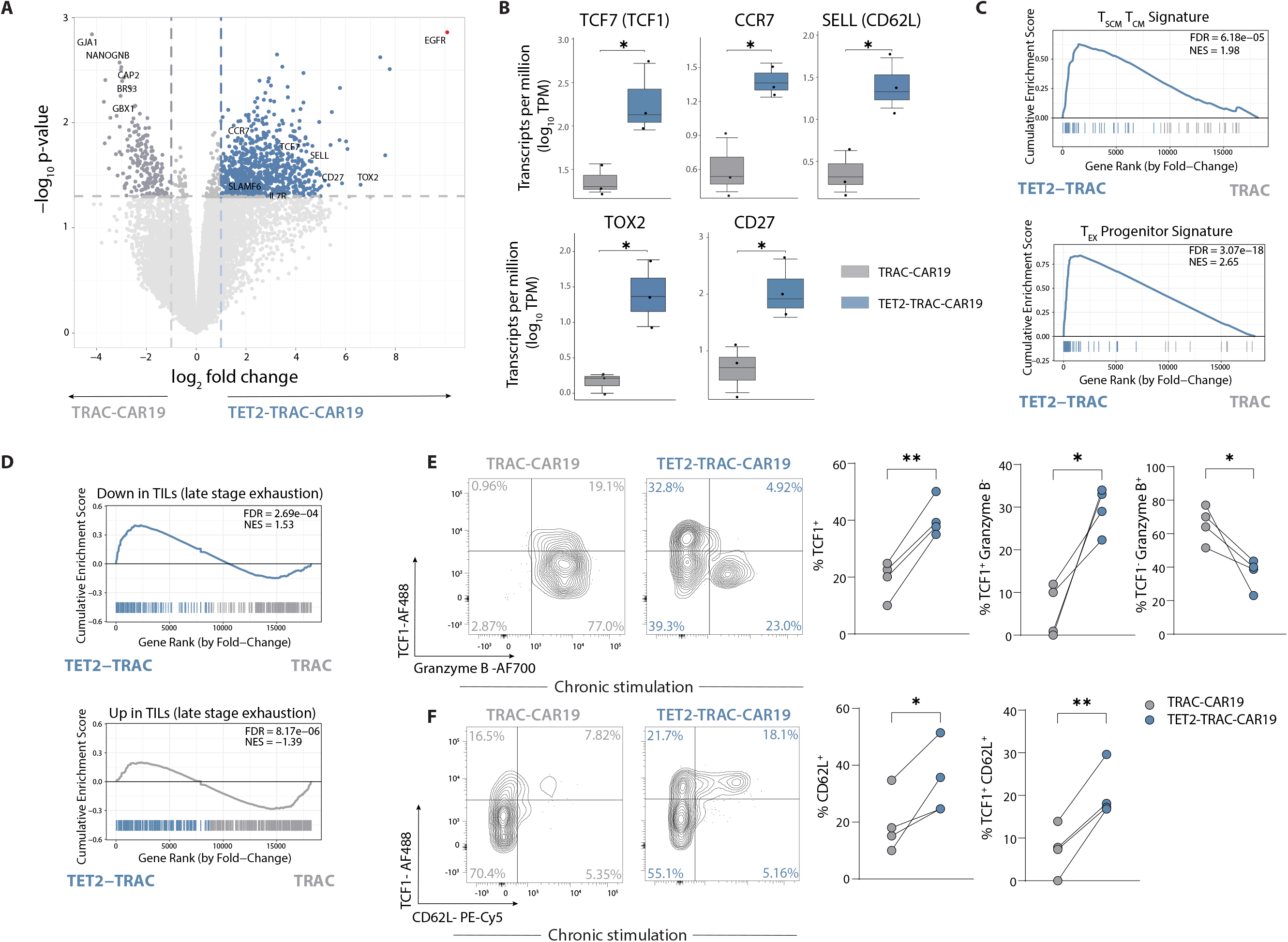
TET2 disruption limits terminal differentiation of TRAC-CAR19 T-cells. (**A**) Volcano plot showing differentially regulated genes in CD8^+^ TET2-TRAC-CAR19 and TRAC-CAR19 T-cells after 4 stimulations; select markers highlighted. EGFR highlighted in red as TET2-KI construct positive control. Graph axes represent log_2_ fold change and -log_10_ p-value, n = 3. (**B**) Box plots of individual gene log_10_ Transcripts per Million (TPM) for TET2-TRAC-CAR19 and TRAC-CAR19 T-cells from (**A**), n = 3. (**C-D**) GSEA for signatures of stem-cell central memory and central memory T-cells (T_SCM,_T_CM_) and T_EX_ progenitors (gene sets from GSE147398) (**C**) and from human tumor infiltrating lymphocytes (TILs) (from *Saleh et al.*). (**D**) Normalized enrichment score (NES) and FDR indicated in panel. (**E**) Example plots and data of CD8^+^ CAR T-cell TCF1/granzyme B subpopulations after 4 stimulations, n = 4. (**F**) Example plots and data of CD8^+^ CAR T-cell TCF1/CD62L subpopulations after 4 stimulations, n = 4. Data shown as mean ± SEM (**B**) or individual values (**E, F**) from independent donors. ns p > 0.05; * p < 0.05; ** p < 0.01; *** p < 0.001 by paired t-test.

### Dual knock-in CAR T-cells display enhanced tumor control in aggressive B-ALL

Finally, we tested the *in vivo* anti-tumor function of CAR T-cells lacking TET2 in an NSG xenograft mouse model for aggressive B-cell ALL (NALM-6) (**Fig. 7A**). NALM-6 cells expressing CD19 and Click Beetle Green luciferase were engrafted into recipient mice and tumor growth was tracked through luminescent imaging. First, we found that *TET2_KO_* CAR T-cells (**fig. S6A**) mediated superior anti-tumor control **(fig. S6B**) and increased survival compared to *AAVS1_KO_* CAR T-cells (**fig S6C**). We next evaluated if TET2-TRAC-CAR knock-in also provided improved tumor control. TRAC-CAR19 T-cells or TET2-TRAC-CAR19 T-cells were administered seven days post-tumor injection. PBS only (i.e., no cells), unedited T-cells (without CRISPR editing or CAR transduction) and TET2-KI only T-cells (without CAR) were administered as controls. All control groups had rapid disease progression and succumbed to ALL by day 31 (**Fig. 7, B-E**), whereas both CAR T-cell experimental groups demonstrated considerable tumor control (**Fig. 7, B, F-G**). TET2-TRAC-CAR19 T-cells showed enhanced efficacy compared to TRAC-CAR19 T-cells **(Fig. 7H)** that was reflected in a substantial reduction in tumor burden by day 32 post-tumor injection (**Fig. 7I**). Sustained tumor control following TET2-TRAC-CAR19 T-cell administration translated into improved survival of mice compared to both the control and TRAC-CAR19 T-cell groups (**Fig. 7J**). Thus, precise *TET2* disruption under the control of a safety switch in allogenic CAR T-cells enhances tumor control and animal survival.

**Figure 7.**
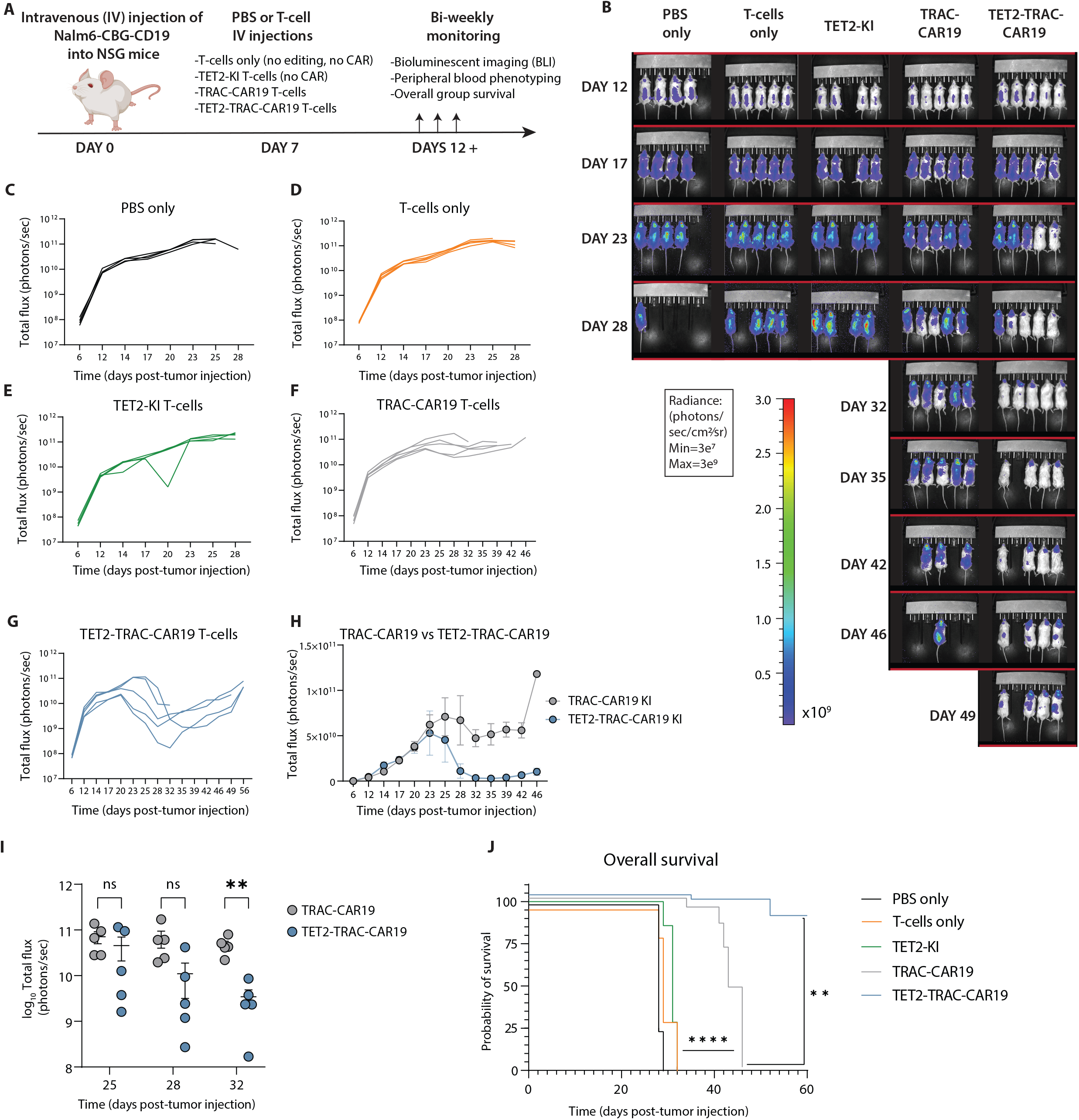
TET2 knock-in enhances the anti-tumor activity of TRAC-CAR19 T-cells *in vivo.* (**A**) Overview of *in vivo* experimental design, n = 4-5 mice per experimental group; one experiment. (**B**) Longitudinal tumor burden of all experimental groups by bioluminescent imaging (BLI). (**C-G**) Tumor outgrowth for PBS only (**C**), T-cell (**D**), TET2-KI T-cell (**E**), TRAC-CAR19 T-cell (**F**) and TET2-TRAC-CAR19 T-cell (**G**) groups, with individual mice shown. (**H**) Longitudinal comparison of TRAC-CAR19 and TET2-TRAC-CAR19 T-cell group tumor burden. (**I**) BLI comparison at days during and immediately after peak CAR T-cell response in TRAC-CAR19 and TET2-TRAC-CAR19 groups, line at mean with SEM; ns p > 0.05; * p < 0.05; ** p < 0.01 by multiple paired t-tests with Holm-Šídák correction. (**J**) Overall group survival, ** p < 0.01; **** p < 0.0001 by Mantel-Cox log-rank test. Data shown as mean ± SEM (**H**) or individual values (**C-G, I**) from each mouse.

## DISCUSSION

We previously reported a case of CAR transgene integration at the *TET2* locus, which shifted T-cell differentiation towards a central memory-like phenotype (*10*). Here, we identified similar insertions among multiple leukemia patients receiving CAR T-cell therapy. The prevalence of these integrations and disruption of *TET2* prompted further investigation into the role of TET2 in regulating T-cell fate and suggested a mechanism through which TET2 could be manipulated to improve CAR T-cell efficacy. Using both *in vitro* and *in vivo* approaches, we identified a role for TET2, a key enzyme in active DNA demethylation, in driving the epigenetic transitions of CD8^+^ T-cells towards terminal differentiation, at the expense of retaining the stem cell-like characteristics of progenitor cells. We took advantage of this function to engineer *TET2*-disrupted CD19 CAR T-cells with improved tumor control. Crucially, the addition of a functional safety switch into this CAR T-cell design provides a therapeutically viable approach to alter T-cell fate for potential patient benefit.

*TET2* deletion restrained terminal differentiation in both chronic LCMV infection *in vivo* and in two *in vitro* CAR T-cell models of exhaustion. In CAR T-cell exhaustion models, this decrease in terminal differentiation resulted in improved proliferative capacity and cytokine production and decreased expression of IRs. A previous study showed that disruption of *TET2* enhanced the *in vivo* expansion of 41BB-costimulated but not CD28-costimulated CAR T-cells (13). However, here, loss of TET2 impacted CAR-T cell phenotype comparably for both 41BBζ- and CD28ζ- CD19 CAR T-cell constructs *in vitro*. The difference in these findings likely reflects differences in the timing of the measured responses and/or context of antigen stimulation. Thus, the role of co-stimulatory domains in TET2-deficient CAR T-cells warrants further investigation, as our data suggest that modulation of TET2 could be used to improve expansion for CAR T-cell products that typically have shorter persistence (*63*).

The decreased terminal differentiation observed in chronically stimulated TET2-deficient CAR T-cells was accompanied by elevated expression of several stem cell and memory-associated genes, including TCF1. Expression of stem cell-like features is a hallmark of the T_EX_ progenitor cells that sustain exhausted CD8^+^ T cell responses during cancer and chronic viral infection (*29–35*). To investigate where TET2 might function in the developmental trajectory of CD8^+^ T cell exhaustion, we turned to the well-characterized LCMV chronic infection model. Indeed, *TET2* deficiency in this setting also protected the T_EX_ progenitor population and restrained development of terminally exhausted T_EX_ cells. Thus, disruption of TET2 enabled maintenance of a progenitor-like population in CAR T-cells and CD8^+^ T cells exposed to chronic viral infection. Furthermore, in chronic viral infection, the terminally exhausted T_EX_ cells that escaped this differentiation block retained key features of T_EX_ progenitors, including decreased expression of several IRs and effector-associated genes. Indeed, *TET2* ablation limited chromatin accessibility for T_EFF_-associated TFs in terminally differentiated T_EX_ cells and reduced expression of TOX. TOX has been proposed to regulate terminal exhaustion (*34*); thus, together these data suggest that TET2 may coordinate with TOX and T_EFF_-associated TFs to initiate and/or maintain the terminal differentiation of T_EX_.

In transitioning to a progenitor-like memory state and away from terminal differentiation, T-cells undergo metabolic reprogramming, including a shift to oxidative metabolism (*64*), that augments proliferation and function. Accordingly, T_CM_ and T_SCM_ cells demonstrate superior anti-tumor potency compared to effector-like cells in CAR T-cell therapies (*10, 65–67*). Metabolic programming also shifts during CD8^+^ T-cell exhaustion. Metabolic fitness, including glucose uptake capacity, may decrease as T_EX_ progenitors increase PD1 expression and terminally differentiate (*18, 68*). Together, these findings highlight commonalities in how metabolic programing is associated with both progenitor-associated biology and terminal differentiation across multiple contexts. *TET2_KO_* CAR T-cells displayed enhanced metabolic fitness post-manufacturing, potentially providing a growth and/or survival advantage. As these metabolic changes were accompanied by an increase in frequency of central memory T-cells, it will be important to disentangle how TET2 may regulate metabolism independently of memory differentiation. However, these data imply first that TET2 could play a critical role in regulating metabolic reprogramming during early T-cell differentiation, and second, that loss of TET2 promotes more progenitor-like metabolism. Therefore, TET2 loss may limit terminal exhaustion, at least in part, through regulation of metabolic function either early in CD8^+^ T-cell differentiation before establishment of exhaustion or as exhaustion progresses.

In settings of both acute and chronic antigen exposure *in vitro* and *in vivo*, TET2 regulated the acquisition (or re-acquisition) of effector-like biology and drove terminal differentiation. These findings suggested that TET2 orchestrates the development of a core effector and terminal differentiation program that coordinates with context dependent TFs and the epigenetic landscape to result in distinct CD8^+^ T-cell fates. Supporting this hypothesis, we also identified a role for TET2 in promoting effector biology and T_EFF_ differentiation before fate-commitment to exhaustion, driving development of short-lived T_EFF_-like cells and away from the formation of the T_EX_ precursor pool (*55*). Thus, TET2 regulates terminal differentiation and the acquisition of effector-like biology at multiple checkpoints in CD8^+^ T-cell differentiation, implying that analogous epigenetic programs are reused throughout CD8^+^ T-cell differentiation. Furthermore, TET2 has been reported to regulate self-renewal and terminal differentiation of additional cell types, including hematopoietic stem cells (*69*). This implies that TET2 may have a broad, cell lineage-independent role in orchestrating terminal differentiation. Further work is needed to understand how TET2 and DNA methylation coordinates these epigenetic transitions and functions with other epigenetic regulators and TFs to regulate cell differentiation in distinct contexts.

TET2 disruption decreased expression of effector-associated molecules, including granzymes, in both chronic stimulation CAR-T cell models and in chronic infection. However, loss of TET2 did not negatively impact CD8^+^ T-cell anti-tumor efficacy. Indeed, TET2-TRAC-CAR19 T-cells exhibited improved tumor control compared to TRAC-CAR19 T-cells. One possible explanation for our findings is that this increased anti-tumor efficacy results from enhanced CD8^+^ T cell proliferation and/or survival, potentially driven by improved maintenance or expansion of the self-renewing stem cell-like progenitor population. Importantly, T_EX_ progenitors retain the ability to produce cytokines, including IL-2, despite low expression of cytotoxic markers (*32*). Thus, strategies that increase the maintenance of progenitor populations, in addition to enhancing effector function, may have potential as a means to improve CAR T-cell efficacy.

Understanding the epigenetic regulation of T_EX_ cell development holds promise for advancing immunotherapies like CAR T-cells. The hyperproliferative phenotype of *TET2*-deficient CAR T-cells underscores the efficacy of epigenetic reprogramming, yet raises substantial long-term safety concerns (*10, 13*). Individual mutations implicated in T-cell lymphoma alone typically do not lead to lymphomagenesis directly; instead, they are often detected in aberrant cells contributing to autoinflammatory or autoimmune disorders (*70*). Additionally, prior studies leveraging gene knockout strategies targeting T-cell lymphoma tumor suppressors have shown no signs of malignant transformation (*71*). However, deliberate disruption of *TET2* for CAR T-cell therapy warrants caution, especially in elderly patients susceptible to acquiring *DNMT3A* mutations (*72*), which can cooperate with *TET2* loss, potentially leading to T-cell oncogenesis (*73*). To address these concerns, we utilized synthetic biology to develop a next-generation cell therapy product. This approach improves the anti-tumor efficacy of CAR T-cells, modulating TET2 to limit terminal exhaustion, while also reducing the risk of lymphomagenesis, autoimmunity, or graft-versus-host disease. First, disrupting *TRAC* mitigates the risk of pathological signaling from the endogenous TCR. Second, our dual knock-in strategy ensures precise insertion of transgenes at specific loci, averting random genomic integration. Third, incorporating a safety switch allows for depletion of *TET2*-disrupted cells if necessary. Additional mitigation strategies could be applied, such as screening for pre-existing mutations predisposing engineered cell products to hyperproliferation or transformation (*74*), administering corticosteroids, which *TET2_KO_* T-cells are highly sensitive to (*10, 13*), and transient or partial suppression of TET2 during CAR T-cell production and/or after infusion. Our findings thus underscore the practical significance and feasibility of targeted epigenetic reprogramming to shape CAR T-cell differentiation, highlighting the potential of TET2 modulation to redirect T_EX_ cell fate.

## MATERIALS AND METHODS

### Study design

The study investigated the role of TET2 in regulating exhausted CD8^+^ T-cell differentiation (T_EX_) in cancer and chronic viral infection, employing human CAR T-cells and a murine lymphocytic choriomeningitis virus (LCMV) model. We evaluated the impact of *TET2* disruption on T_EX_ phenotypes, differentiation fate, and functional outcomes both *in vitro* and *in vivo*. Flow cytometry assessed protein expression related to T-cell exhaustion and memory differentiation, while ATAC-seq explored chromatin accessibility landscapes across T_EX_ subsets. Transcriptomic analysis elucidated underlying pathways and transcriptional regulation by TET2. Additionally, we developed a novel CRISPR/Cas9-based genome editing approach to engineer allogeneic CAR T-cells for enhanced anti-tumor responses through TET2 modulation of T_EX_. Sample sizes were estimated based on preliminary experiments, with *in vitro* functional assays performed at least three times. Investigators were not blinded during experiments or outcome assessment.

### Primary human cells

CAR T-cells and control samples were generated from healthy donor peripheral blood mononuclear cells (PBMCs) through leukapheresis, following University of Pennsylvania Institutional Review Board-approved protocols. Written informed consent was obtained from all participants, consistent with the principles outlined in the Declaration of Helsinki, International Conference on Harmonization Guidelines for Good Clinical Practice, and the U.S. Common Rule.

### Cell lines

For viral vector production, HEK 293T cells and GP2-293, a HEK 293-derived retroviral packaging cell line, were cultured in hR10 medium (RPMI 1640 supplemented with 10% heat-inactivated FBS, 2% Hepes buffer, 1% GlutaMAX, and 1% penicillin-streptomycin). SUP-T1 were used to determine lentiviral titers. HEK 293T cells were sourced from the American Type Culture Collection (ATCC) and GP2-293 from Takara Bio. NALM-6 cells expressing click beetle green luciferase and green fluorescent protein (CBG-GFP), provided by Marco Ruella at the University of Pennsylvania, were utilized. K562 human leukemia cell lines including a variant expressing the extracellular domain of the CD19 protein were obtained from Carl H. June at the University of Pennsylvania and maintained in hR10 medium. 143B osteosarcoma cells modified to express GFP and firefly luciferase, along with NALM-6 cells engineered for GD2 expression, were also cultured in hR10 medium. Cell line authenticity was confirmed via short-tandem-repeat profiling meeting the International Cell Line Authentication Committee’s guidelines with more than 80% match, conducted by the University of Arizona Genetics Core. Regular mycoplasma screenings were performed to ensure cell line health and purity before and after genetic modifications.

### Analysis of *TET2* integration sites in CAR T-cell-treated leukemia patients

Genomic DNA from patient cell sources (whole blood, bone marrow, PBMCs, or T-cells; pre- and post-infusion) was isolated for library preparation followed by paired-end Illumina sequencing, as previously described (*10, 75*). CLL and ALL human integration site data were aligned to the human genome hg38 and analyzed using a previously published integration analysis pipeline (*76*). Sample timepoints were grouped into four categories (day 0, days 1-15, days 16-31, day 31+). Percent relative abundance represents the estimated proportion of cells with integration in a sample.

### Lentiviral packaging

Briefly, HEK 293T cells were transfected with 7 μg of pVSV-G glycoprotein envelope plasmid, 18 μg of pMDLg/p.RRE Gag/Pol plasmid and 18 μg of pRSV.Rev plasmid alongside 15 μg of transfer vector plasmid encoding for CAR of interest using Lipofectamine 2000 (Thermo Fisher Scientific) and Opti-MEM (Gibco). Cell culture supernatant was harvested 24- and 48-hours post-transfection, centrifuged at 900 RCF for 10 minutes at 4°C and filtered through a 0.45 μM vacuum filter. Following filtration, 24-hour supernatant was concentrated by ultracentrifugation at 8877 RCF overnight at 4°C, while 48-hour supernatant was concentrated overtop of the overnight viral pellet at 76,790 RCF for 2 hours at 4°C. Concentrated virus was stored at −80°C.

### T-cell culture and lentiviral transduction

T-cells were isolated from healthy donor PBMCs using the Pan T-cell Isolation Kit following manufacturer’s instructions (Miltenyi Biotec). Isolated T-cells were activated using anti-CD3/CD28 antibody-coated Dynabeads (Thermo Fisher Scientific) at a 3:1 bead-to-cell ratio in T-cell media consisting of OpTmizer CTS SFM media (Thermo Fisher Scientific) supplemented with 5% human AB serum and 100 Units/mL human IL-2 (PeproTech). After a 24-hour incubation, lentivirus containing the appropriate CAR construct was introduced to the culture at a multiplicity of infection (MOI) of 2.5. CAR T-cell expansion proceeded following established protocols (*77*).

### AAV construct design

DNA sequences containing either a truncated EGFR (tEGFR) sequence driven by an EF1α promoter (for TET2-KI), or a truncated NGFR (tNGFR) sequence, T2A sequence and an anti-CD19 single-chain variable fragment (scFv) fused to 4-1BB and CD3ζ stimulatory endodomains (for TRAC-CAR19-KI) were subcloned into recombinant AAV6 plasmids (GenScript). DNA sequences were flanked with 400 base-pair homology arms immediately upstream and downstream of the TET2 gRNA or TRAC gRNA cut sites, respectfully. Large-scale packaging of AAV6 virus was done by co-transfection of a packaging cell line with the rAAV6 transgene plasmid of interest, a rep- and cap-encoding plasmid and an adenovirus-derived replication helper plasmid (Charles River).

### CRISPR-Cas9–mediated editing and AAV transduction

*TET2* and *TRAC* editing via CRISPR-Cas9 was conducted 72 hours post-T-cell activation. Single guide RNA (sgRNA) reagents from Integrated DNA Technologies targeted the *TET2* and *TRAC* loci. The sgRNA sequences with protospacer-adjacent motif (PAM) sequences are indicated as follows: *TET2* 5’-<u>CGG</u>GGATACCTATACAGATCCAT-3’ and *TRAC* 5’-<u>AGG</u>GAGAATCAAAATCGGTGAAT-3’. The control *AAVS1* targeted sequence is: 5’-CCATCGTAAGCAAACCTTAG<u>AGG</u>-3’.

Activated T-cells were de-beaded magnetically, washed with 1X PBS at 300 × g for 5 minutes, and resuspended in P3 4D-nucleofection buffer (Lonza). TrueCut Cas9 Protein v2 (Thermo Fisher Scientific). sgRNAs targeting *TET2* and/or *TRAC* were individually complexed at 6µg:3.2µg for 10 minutes at room temperature to form ribonucleoprotein (RNP) complexes before nucleofection. Nucleofection into T-cells was performed using a Lonza 4D Nucleofector X Unit with high fidelity program EO-115, followed by a 10-minute resting period. For AAV-mediated knock-in, cells were transduced with AAV viral vectors carrying TRAC-CAR19-tNGFR and/or TET2-tEGFR constructs (Charles River) at an MOI of 50,000.

*TET2* knockout efficiency was confirmed by isolating genomic DNA from CAR T-cells at Day 7 using the dNeasy Blood & Tissue Kit (Qiagen). PCR of genomic DNA was performed with *TET2* Forward Primer 5’-TCCCTGAGTCCCAGTCCATC-3’ and Reverse Primer 5’-TCAGGAATGGCCAGGTTCTG-3’ using MyTaq Red 2X Mix (Meridian Bioscience). Purified control and edited PCR products underwent Sanger sequencing (Azenta), and editing efficiency was determined by Tracking of Indels by DEcomposition (TIDE) through comparison of control and edited Sanger sequence electropherogram files.

To confirm tEGFR and tNGFR-CAR19 construct knock-ins, genomic DNA was isolated from end-of-expansion transduced CAR T-cells. PCR of genomic DNA was carried out with the following primer sets: *TET2* (unedited) Forward Primer 5’-TCCCTGAGTCCCAGTCCATC-3’, *TET2* (unedited) Reverse Primer 5’-TCAGGAATGGCCAGGTTCTG-3’, *TET2* (edited) Forward Primer 5’-CATCACGAGCAGCTGGTTTC-3’, *TET2* (edited) Reverse Primer 5’-GGCAATTGAACCGGTGCCTA-3’, *TRAC* (unedited) Forward Primer 5’-TCCCTGAGTCCCAGTCCATC-3’, *TRAC* (unedited) Reverse Primer 5’-CTTCATGCCCTGCATCTCCA-3’, *TRAC* (edited) Forward Primer 5’-CATCACGAGCAGCTGGTTTC-3’, *TRAC* (edited) Reverse Primer 5’-CATCAGTTGCAGGGCAAGTC-3’. Edited and unedited PCR products underwent purification 648 and Sanger sequencing.

### Western blot analysis of *TET2* knockout

CAR T-cells were lysed in 1X lysis buffer (Cell Signaling Technology) and supernatants were collected after centrifugation. Cell lysate samples (30μg) were separated on a NuPAGE 4-12% Bis-Tris gel (Invitrogen) and transferred onto a membrane using the iBlot 2 Dry Blotting System (Invitrogen). The membrane was blocked with 5% skim milk and probed with primary antibodies overnight at 4°C: either rabbit monoclonal anti-TET2 (Cell Signaling Technology) or monoclonal mouse anti-GAPDH (Thermo Fisher). Primary antibodies were diluted in 1X PBS with 0.2% Tween and 5% BSA. After washing, the membrane was incubated with goat anti-mouse or anti-rabbit HRP-linked secondary antibody (Thermo Fisher) for 1 hour at room temperature. Finally, the membrane was treated with equal parts of Pierce ECL Western blotting substrate (Thermo Fisher Scientific) and visualized.

### Flow cytometry of human immune cells

Cells were collected and stained with LIVE/DEAD Fixable Aqua Dead Cell Stain Kit (Invitrogen) for 20 minutes at room temperature. After washing with hFACS buffer (PBS + 2% FBS + 0.05% Sodium Azide), surface antibodies were incubated with cells for 30 minutes at 4°C in hFACS buffer and Brilliant Stain Buffer (BD Biosciences). For intracellular staining, samples were fixed and permeabilized using the FoxP3 Transcription Factor Staining Buffer Kit (Thermo Fisher Scientific) for 30 minutes, followed by staining with intracellular antibodies for an additional 30 minutes. Data acquisition was performed using a BD LSRFortessa and analyzed with FlowJo^TM^ software (BD Life Sciences). Compensation setup utilized Anti-Mouse Ig, κ and Anti-Rat/Hamster Ig, κ CompBeads (BD Biosciences) along with Fluorescence Minus One (FMO) controls to establish gating boundaries. SPICE plots were generated from single gated inhibitory receptors, grouped using Boolean ‘AND’ gates, and plotted using SPICE 6.1 software (https://niaid.github.io/spice/). Refer to **table S6** for antibody details.

### Seahorse metabolic flux assay

Using a Seahorse xFe96 Analyzer (Agilent), we conducted the Seahorse Mitochondrial Stress Test. The xFe96 Pro sensor cartridge (Agilent) was hydrated overnight with sterile water at 37°C in a non-CO_2_ incubator, followed by XF Calibrant (Agilent) hydration. Thawed T-cells were rested overnight, washed with 1X PBS, and treated with Seahorse Assay Media. Cells (1-2 × 10^5^) were plated in Poly-D-Lysine coated 96-well microplates (Agilent) with 4-5 technical replicates. The microplate was centrifuged and incubated to facilitate cell attachment. Drug solutions (oligomycin, FCCP, Rotenone/Antimycin A) were prepared in the sensor cartridge. The assay measured basal OCR and ECAR triplicate at baseline and after each drug addition using WAVE software (Agilent).

### CAR T-cell serial restimulation assay

CAR^+^ T-cells were purified using a biotin-conjugated AffiniPure Goat Anti-Mouse IgG F(ab’)₂ fragment specific antibody and anti-biotin microbeads (Miltenyi Biotec). In the case of dual knock-in TET2-TRAC CAR T-cells, EGFR^+^, NGFR^+^, and EGFR/NGFR-dual positive cells were purified using PE and APC-conjugated antibodies alongside anti-PE, anti-APC, or anti-PE MultiSort microbeads as per the manufacturer’s instructions (Miltenyi Biotec). Purity was assessed by flow staining against knock-in markers tEGFR and tNGFR. K562-CD19^+^ cells were exposed to 100 Gy ionizing radiation using the xRad320 (Precision X-Ray). CAR T-cells were co-cultured with irradiated K562-CD19^+^ cells at a 1:1 ratio with 1 million CAR T-cells per 1 million K562 cells in hR10. Co-culture supernatants were harvested 24 hours after each stimulation and frozen at −20°C. At 5 days post-stimulation (defined as acute stimulation), absolute CAR T-cell counts were assessed with a LUNA-FL Dual Fluorescence Cell Counter (Logos Biosystems) and re-cultured at a 1:1 ratio with fresh hR10 media and newly irradiated K562 cells for 4-5 additional stimulations (defined as chronic stimulation). CAR T-cells were then cryopreserved for phenotyping and transcriptomic profiling.

### Cytokine analysis

Supernatant cytokines were quantified flow cytometrically using the LEGENDplex™ Human CD8/NK Panel as per the manufacturer’s instructions (BioLegend). Data was acquired on LSRFortessa and data analysis was performed with BioLegend LEGENDplex™ Data Analysis Software Suites (BioLegend Qognit Cloud Platform).

### Assay for exhaustion in CAR T-cells with high tonic signaling

CAR T-cells were transduced with retrovirus on days 2 and 3 post-activation. Briefly, 12-or 24-well plates, non-tissue-culture-treated, were coated with 1 mL or 500 µL, respectively, of 25 μg/mL Retronectin (Takara) in PBS and incubated at 4°C overnight. The following day, plates were washed with PBS and then blocked with 2% BSA in PBS for 10 minutes. Retroviral supernatants were added, and plates were centrifuged at 32°C for 2 hours at 2500 RCF. After centrifugation, viral supernatants were removed, and T-cells were seeded into each virus-coated well at a density of 1 × 10^6^ T-cells/well for 12-well plates and 0.5 × 10^6^ T-cells/well for 24-well plates. CRISPR knockout of *TET2* (or *AAVS1* as a control) was performed 2-4 days post T-cell activation to achieve maximal editing efficiency, using the EH115 program on a Lonza 4D Nucleofector. Cells were immediately recovered in 260 µL of warm complete AIM-V media supplemented with 500 U/mL IL-2 in round-bottom 96-well plates and expanded into 1 mL fresh medium after 24 hours. Cells were maintained at densities of 0.5-2 × 10^6^ cells per mL in well plates until day 14-16 for functional and phenotypic characterization. Editing efficiency was assessed using TIDE as described previously. Immunophenotyping of CAR T-cells via flow cytometry was performed on days 11 and 15 of expansion. Cytotoxicity of HA.28ζ CAR T-cells was evaluated using an Incucyte^®^ Live-Cell Analysis System at day 15 at the end of expansion. In brief, 25 × 10^5^ GFP^+^ 143b-GL osteosarcoma tumor cells were seeded in triplicate in 96-well plates and co-cultured with T-cells at effector:target ratios of 1:1, 1:2, 1:4, 1:8, and/or 1:16 in 300 µL of T-cell medium without IL-2 in 96-well flat-bottom plates. Plates were imaged at 10X zoom with 4-9 images per well every 2-4 hours for 96 hours using the IncuCyte ZOOM Live-Cell analysis system. Total integrated GFP intensity per well or total GFP area (µm^2^/well) were used to analyze expansion or contraction of 143B cells, with four images captured per well at each time point. Total tumor GFP fluorescence (normalized to the initial *t* = 0 timepoint) was recorded, and the normalized tumor GFP signal was used as the cytolysis threshold.

Cell culture supernatants from 1:1 E:T co-cultures were utilized to determine IL-2 and IFN-γ concentrations via ELISA. Specifically, 5 × 10^4^ CAR T-cells were co-cultured with 5 × 10^4^ tumor cells in 200 µL of complete T-cell medium (AIM-V or RPMI) without IL-2 in a 96-well plate, all in triplicate. After 24 hours of coculture, culture supernatants were collected, diluted 20 to 100-fold, and analyzed for IL-2 and IFN-γ using ELISA MAX kits and Nunc Maxisorp 96-well ELISA plates. Absorbance readings were obtained using a Spark plate reader (Tecan Life Sciences).

### LCMV mouse studies

Mice were maintained in a specific-pathogen-free facility at the University of Pennsylvania, in accordance with the Institutional Animal Care and Use Committee. B6;129S-*Tet2^tm1.1Iaai/^J* (TET2^fl/fl^) mice and CD4^Cre+^ mice were obtained from the Jackson Laboratory (JAX). TCR transgenic P14 C57BL/6 mice expressing a TCR specific for LCMV peptide D^b^GP^33–41^ (*78, 79*) were bred in house. All mice were backcrossed to and maintained on a C57BL/6J background. P14 mice were bred to TET2^+/+^ CD4^Cre+^ or TET2^fl/fl^ CD4^Cre+^ mice to generate WT (TET2^+/+^ CD4^Cre+^ P14^+^) and *TET2*_KO_ (TET2^fl/fl^ CD4^Cre+^ P14^+^) P14 donor mice. For all experiments, WT and *TET2_KO_* donor mice were age and sex matched. For P14 co-transfer experiments, sex-matched recipient C57BL/6 mice were purchased from JAX at 5-8 weeks of age.

### Chronic LCMV infection

Recipient mice were infected intravenously (i.v.) with 4 × 10^6^ PFU of LCMV clone 13. LCMV titers were determined via plaque assay as described (*80*).

### Naïve P14 cell co-transfer

Adoptive transfer of P14 cells was performed as described (*8*). P14 cells were isolated from the peripheral blood of naïve congenically distinct WT and *TET2_KO_* donor mice using a histopaque 1083 gradient (Sigma-Aldrich). WT and *TET2_KO_* P14 cells were mixed at a 1:1 ratio and a total of 500 P14 cells (250 WT and 250 *TET2_KO_*) were adoptively transferred intravenously into recipient mice of a third congenic background. The 1:1 ratio was confirmed by flow cytometry (BD LSRII). One day post-adoptive transfer, recipient mice were infected with LCMV clone 13 (day 0). Unless otherwise indicated, recipient mice were treated with CD4-depleting antibody (GK1.5, 200 mg/injection) on day −1 and day +1 relative to infection with LCMV clone 13.

### Retroviral transduction of the TET2 catalytic domain

The FLAG-tagged murine TET2 catalytic domain in pMXs was provided by R. Kohli (University of Pennsylvania) and subsequently inserted into MIGR1 courtesy of Warren Pear (University of Pennsylvania), with an expanded multiple cloning site introduced. Empty MIGR plasmid was used as a control. Retroviruses (RV) were generated in HEK 293T cells. P14 cells from either WT or *TET2_KO_* donor mice were activated, and retroviral transduction performed as previously described (*81, 82*). CD8^+^ T-cells were isolated from spleens of P14 donor mice by negative selection using the EasySep^TM^ Mouse CD8^+^ T-cell isolation kit (STEMCELL Technologies). P14 cells were activated *in vitro* for 24-28 hours with 100U/mL recombinant IL-2, 1 mg/mL LEAF anti-mouse CD3e and 0.5 mg/mL LEAF anti-mouse CD28 in mouse R10 media (mR10: RPMI-1640 supplemented with 10% FCS, 50U/mL penicillin and streptomycin, l-glutamine, 20mM HEPES, non-essential amino acids (1:100), 1mM sodium pyruvate, and 50mM b-mercaptoethanol). Activated P14 cells were transduced by spinfection at 2,000 x g for 90 min at 32°C in mR10 + 100U/mL IL-2 and 0.5 mg/mL polybrene. WT P14 cells were transduced with MIGR1, while *TET2_KO_* P14 cells from donor mice of a distinct congenic were transduced with either MIGR1 or TET2 CD. After 24 hours rest, P14 cells expressing the retroviral reporter GFP were sorted (BD FACS Aria, 37°C) and WT and *TET2_KO_* P14 cells mixed in a 1:1 ratio before adoptive transfer i.v. into recipient mice of a third congenic background. The 1:1 ratio was confirmed by flow cytometry (BD LSRII). 4-5 × 10^4^ total P14 cells were transferred per LCMV clone 13-infected recipient mouse. Recipient mice were infected with LCMV clone 13 on the same day as P14 cell activation.

### Peptide Stimulation, flow cytometry, and sorting of murine immune cells

PBMC were isolated from peripheral blood by repeated lysis with Ammonium-Chloride-Potassium (ACK) lysis buffer and immediately stained in mouse FACS Buffer (mFACS Buffer; PBS + 3% FCS + 2mM EDTA). Splenocytes were processed to a single cell suspension by mechanical disruption over a 70 µm filter, followed by ACK lysis and then counted.

Splentocyte samples were either aliquoted in mFACS Buffer for staining or resuspended in mR10 and stimulated *ex vivo* with LCMV peptide D^b^GP^33^ (0.2 μg/mL) in the presence of Golgi Plug and Golgi Stop (BD Bioscience) for 5 hours at 37°C. Following stimulation, samples were washed in PBS, incubated with a viability dye (15 minutes at RT), and stained with an antibody cocktail targeting surface markers in mFACS Buffer + Brilliant Stain buffer for 30 min at 4°C or 1 hour at RT. In some panels, samples were stained with gp33 tetramer for 1 hour at 37°C in mR10 prior to surface staining. Biotinylated primary antibodies were detected with streptavidin-conjugated secondary antibody for 30 minutes at 4°C. For intracellular staining, samples were permeabilized using the eBioscience^TM^ FoxP3 Transcription Factor Staining Buffer Kit or BD Cytofix/Cytoperm^TM^ Fixation/Permeabilization kit and incubated with intracellular antibodies for 30 minutes at 4°C or 1 hour at RT then washed and stored in BD Stabilizing Fixative until acquisition. Samples were acquired on a BD LSRII or a BD FACSymphony A5 and analyzed in FlowJo^TM^ v10.8 software. Voltages on flow cytometry machines were standardized using fluorescent targets and Spherotech rainbow beads.

For cell sorting for sequencing, splenocytes were stained with a surface antibody cocktail in mR10 + Brilliant Stain Buffer for 30 minutes at 4°C. Samples were sorted on a BD FACSAria at 4°C into mR10 media with 50% FCS. Sorting accuracy was confirmed through post-sort purity checks. See **table S7** for antibody information.

### Bulk RNA-sequencing

For *in vivo* LCMV sample preparation, WT and *TET2_KO_* P14 cells were co-transferred into recipient mice as described in “**Naïve P14 cell co-transfer**”. Total WT and *TET2_KO_* P14 cells and T_EX_ subsets were isolated at day 15 p.i. with LCMV clone 13 with >95% purity, as detailed above. 2 × 10^4^ cells were sorted in triplicate per sample and stored at −80°C in RLT buffer (Qiagen). RNA was isolated using the Qiagen RNeasy Micro Kit, and cDNA libraries were generated following the manufacturer’s instructions with the SMART-Seq v4 Ultra Low Input RNA Kit and Nextera XT DNA library kit. After quantification with the KAPA Library Quant Kit, cDNA libraries were pooled and diluted to 1.8 pg/mL and paired-end sequencing was conducted on a NextSeq 550 (Illumina) using a NextSeq 500/550 Mid Output Kit v2.5 (150 cycles).

For *in vitro* CAR T-cell sample preparation, CD8^+^ T-cells were enriched via positive selection (Miltenyi Biotec), resuspended in TRIzol™ (Thermo Fisher Scientific), and stored at - 80°C. Upon thawing, total RNA was extracted, treated with Dnase, and further processed using an RNA Clean and Concentrator Kit (Zymo Research). Bulk RNA sequencing was performed by Novogene using the NovaSeq6000 system with a paired-end 150bp approach, generating 6 GB of sequencing read data per sample.

Mouse reads were aligned to transcriptome mm39 using STAR with quantification via cufflinks, while human reads were pseudoaligned to the human transcriptome GRCh38 using kallisto. Data was imported into R, transformed into log2 counts per million, and normalized using Trimmed Mean of M-values (TMM) with EdgeR. Differential Gene Expression was determined with linear modeling and adjusted p-values using the Benjamini-Hochberg correction method with limma and EdgeR.

Gene Set Enrichment Analysis (GSEA) was conducted using GSEA software from UC San Diego and Broad Institute developers. Filtered, normalized expression data was utilized as input, with parameters including a weighted enrichment statistic and Signal2Noise metric for gene rankings.

### ATAC-sequencing

For each LCMV T_EX_ sample, 1-2 × 10^4^ P14 cells were sorted in duplicate or triplicate and processed as previously described (*83*) with minor modifications (*82*). Briefly, P14 cells were washed with cold PBS, resuspended in 50 μL of cold lysis buffer (10 mM Tris-HCl, pH 7.4, 10 mM NaCl, 3 mM MgCl_2_, 0.1% IGEPAL CA-630), and centrifuged (750 x g, 10 minutes, 4°C) to remove lysates. Nuclei were immediately resuspended in 25 μL of the transposition reaction mix (12.5μL 2x TD Buffer (Illumina), 1.25μL Tn5 Transposases, 11.25μL nuclease-free H_2_O) and incubated at 37°C for 45 minutes. Transposed DNA fragments were purified using the QIAGEN Reaction MiniElute Kit, barcoded with NEXTERA dual indexes (Illumina), and PCR amplified with NEBNext High Fidelity 2x PCR Master Mix (New England Biolabs). Following purification with the PCR Purification Kit (QIAGEN), fragment sizes were confirmed using the 2200 TapeStation and High Sensitivity D1000 ScreenTapes (Agilent). ATAC-sequencing libraries were quantified, pooled, and sequenced as described above for RNA sequencing. Alignment to the mm39 genome was performed using bwa-mem, and peak calling was performed using MACS2. Differential peak analysis was performed using limma-voom, and motif enrichment was performed using HOMER.

### Antibody-dependent cellular cytotoxicity (ADCC) co-culture

*TET2*-KI T-cells were rested overnight in hR10 media, while donor-matched natural killer (NK) cells were isolated using an NK cell isolation kit (Miltenyi Biotec) and cultured overnight in hR10 supplemented with IL-15 at 10 ng/mL (PeproTech). The following day, TET2 knock-in T-cells were incubated with a Cetuximab biosimilar (R&D Systems, #MAB9577) at a concentration of 2000 ng/mL for 20 minutes. T-cells were then co-cultured at a 1:10 ratio with NK cells, with T-cell numbers normalized to EGFR^+^ expression. After 16 hours, co-cultures were harvested and stained with LIVE-DEAD Aqua, CD3, CD56, and a human IgG PE-conjugated secondary antibody. EGFR expression on *TET2*-KI T-cells alone compared to NK co-culture with or without Cetuximab (gated on live, CD56^−^ CD3^+^ EGFR^+^) was used to calculate percent EGFR decrease as a readout for ADCC.

### Mouse xenograft studies

Male NOD/SCID/IL-2Rγ-null (NSG) mice, aged 7 weeks, were utilized for xenograft studies. Mice received an intravenous injection of 3 × 10^5^ NALM-6 tumor cells expressing a CBG luciferase reporter suspended in 200 µl of PBS. Tumor engraftment was confirmed on day 6 via intraperitoneal injection of IVISbrite D-Luciferin Potassium Salt Bioluminescent Substrate (XenoLight, PerkinElmer), followed by bioluminescent imaging (BLI) using the IVIS® Lumina III In Vivo Imaging System (PerkinElmer). On day 7, mice were administered with 5 × 10^5^ control T-cells, experimental CAR T-cells, or PBS alone. Biweekly tumor imaging was conducted using the IVIS Lumina system after IP luciferin injection to monitor tumor growth or reduction. Regions of interest (ROIs) were delineated around mice for the calculation of bioluminescent tumor burden. Peripheral blood samples were collected via cheek bleeding at the peak of CAR T-cell expansion and lysed with ACK Lysing Buffer (Gibco) to obtain T-cells for immunophenotyping via flow cytometry. Absolute cell counts were determined using 123count eBeads™ Counting Beads (Invitrogen). Kaplan-Meier survival curves were generated based on endpoint survival data.

### Statistical analyses

Summary data are presented as mean ± SEM, as indicated in the figure legends, alongside corresponding p-values. Pairwise sample comparisons were evaluated using a paired *t*-test. For multiple pairwise comparisons, multiple paired *t*-tests with Holm-Šídák correction were used. One-way ANOVA analysis was used for comparisons involving multiple groups, initially assessing differences in mean values with a global omnibus F-test, followed by post-hoc analysis for multiple comparisons if the initial test yielded significance (p < 0.05). Mouse survival was analyzed using the Mantel-Cox log-rank test. Number of donors, animals and experiments are indicated in the figure legends. All statistical analyses were conducted using Prism 9 or 10 (GraphPad Software), and significance was defined as p < 0.05.

## Acknowledgements

The authors are grateful to the Human Immunology Core at the University of Pennsylvania (RRID SCR_022380) and the Hospital of the University of Pennsylvania Apheresis Unit for their provision of peripheral blood mononuclear cells. We acknowledge the Stem Cell and Xenograft Core at the University of Pennsylvania (RRID SCR_010035) for their husbandry services and support with *in vivo* mouse studies, the Cell and Animal Radiation Core at the University of Pennsylvania (RRID SCR_022377) for access to the xRad irradiator, and the Penn Cytomics and Cell Sorting Resource Laboratory (RRID SCR_022376). We thank Dr. Fengjuan Zhang from the Translational and Correlative Sciences Laboratory (TCSL) for helpful advice regarding gene set enrichment analysis and acknowledge the contributions of the TCSL and the Product Development Laboratory (PDL) from the University of Pennsylvania Center for Cellular Immunotherapies. Special thanks to Dr. Megan Davis (Director of PDL) and Dr. Kathrine Alexander from the University of Pennsylvania Epigenetics Institute for their invaluable discussions. We also extend our gratitude to Dr. Carl June (University of Pennsylvania) for his insightful advice and input on translational aspects of TET2 modulation in T-cell therapy. Schematic illustrations were created with BioRender.

## Author contributions

A.J.D., A.E.B., M.S.J., and J.A.F. designed the study. A.J.D., A.E.B., J.R.G., I-Y.J., R.O., G.V., J.K.J., R.M.Y., J.J.M., S.L.M, B.L.L., N.V.F., S.L.B., S.A.G., D.L.P., F.H., M.H.P., F.D.B., E.W.W., E.J.W., M.S.J., and J.A.F. developed methodologies and provided resources, technical feedback as well as intellectual input. A.J.D., A.E.B., C.R.H., C.H.H., G.T.R., W.K., V.W., R.B., S.D., K.D., C.R.H., O.M.K., Z.C., A.C. and N.G. performed experiments. G.M.C., H.H., and S.C. performed RNA- and ATAC-sequencing data analysis. J.K.E., K.A. and S. S-M. performed integration site analyses. The paper was written by A.J.D., A.E.B., M.S.J., and J.A.F., with all authors contributing to writing and providing feedback.

## Data and materials availability

All study data are available within the paper and its supplementary materials. Sequencing data are deposited in Gene Expression Omnibus (GEO) under accession number GSE261093 and integration site sequencing data in the Sequence Read Archive (SRA BioProject PRJNA1093497), both hosted by the National Center for Biotechnology Information (NCBI). Visualization code for TET2 patient integration site data analysis is available at https://github.com/helixscript/TET2_ALL_CLL. For additional data or material inquiries, contact the Penn Center for Innovation at. Requests will be promptly reviewed for any intellectual property or confidentiality issues, and eligible data and materials will be shared following a material transfer agreement. For further assistance, contact the corresponding authors.

---

**Supplemental Figure 1.**
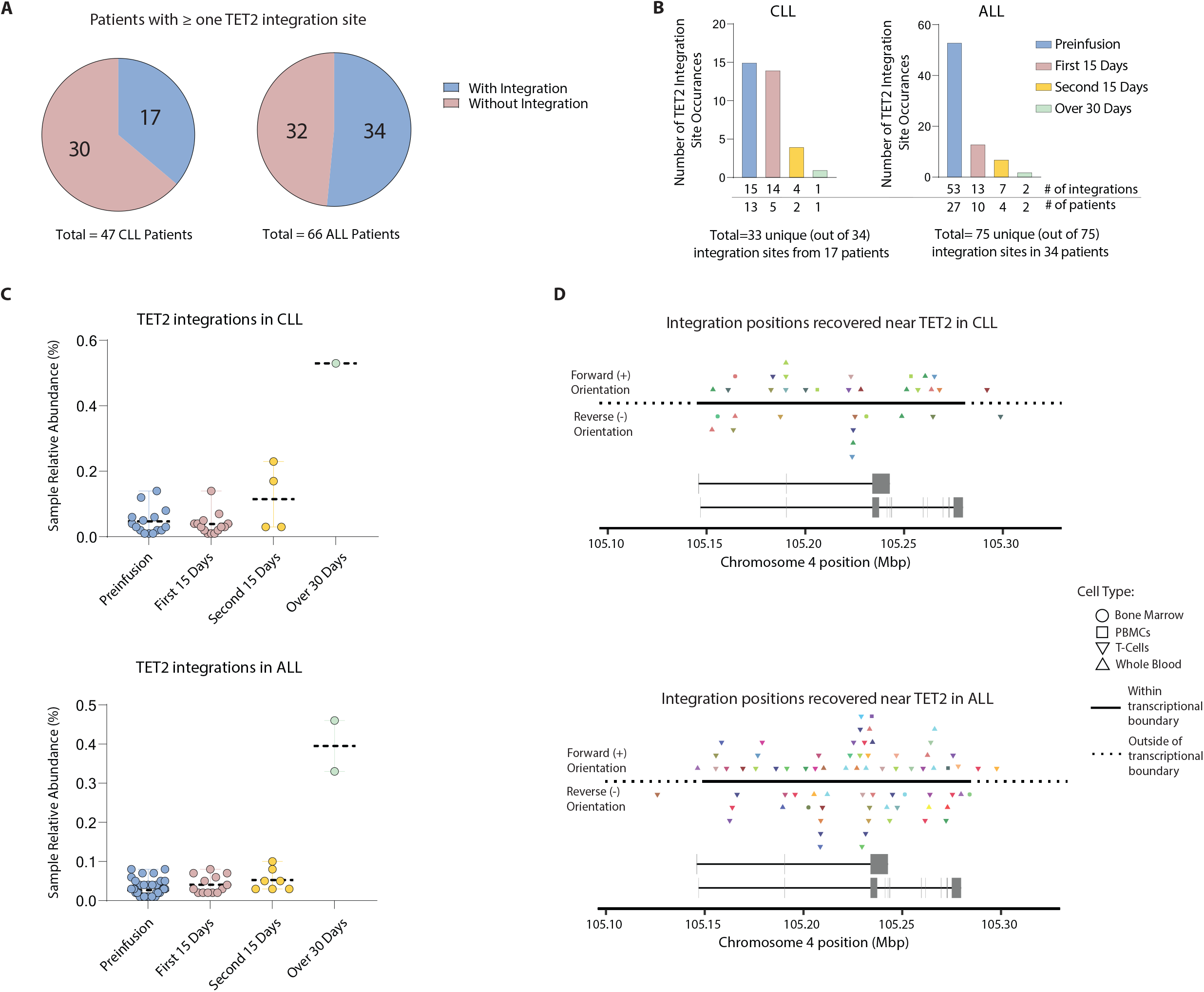
The TET2 locus is a frequent site of lentivirus integration in CAR T-cell treated patients. (**A**) Number of chronic lymphocytic leukemia (CLL) and acute lymphocytic leukemia (ALL) patients with or without ≥ one observed CAR lentiviral integration into *TET2.* (**B**) Distribution of the number of *TET2* integration sites at specified binned timepoints. For CLL patient p04409-09, the same integration was detected within the first 15 days and again in the second 15 days. (**C**) Scatter plots showing the relative abundance of each *TET2* patient integration at specified timepoints for CLL (**top**) and ALL (**bottom**) patient cohorts. Each dot represents one integration site. Dotted line represents mean, error bars represent SEM. (**D**) Location of each integration clone within the *TET2* transcriptional boundary for CLL (**top**) and ALL (**bottom**) patient cohorts. Color coded for patient; symbol shape represents cell type profiled.

**Supplemental Figure 2:**
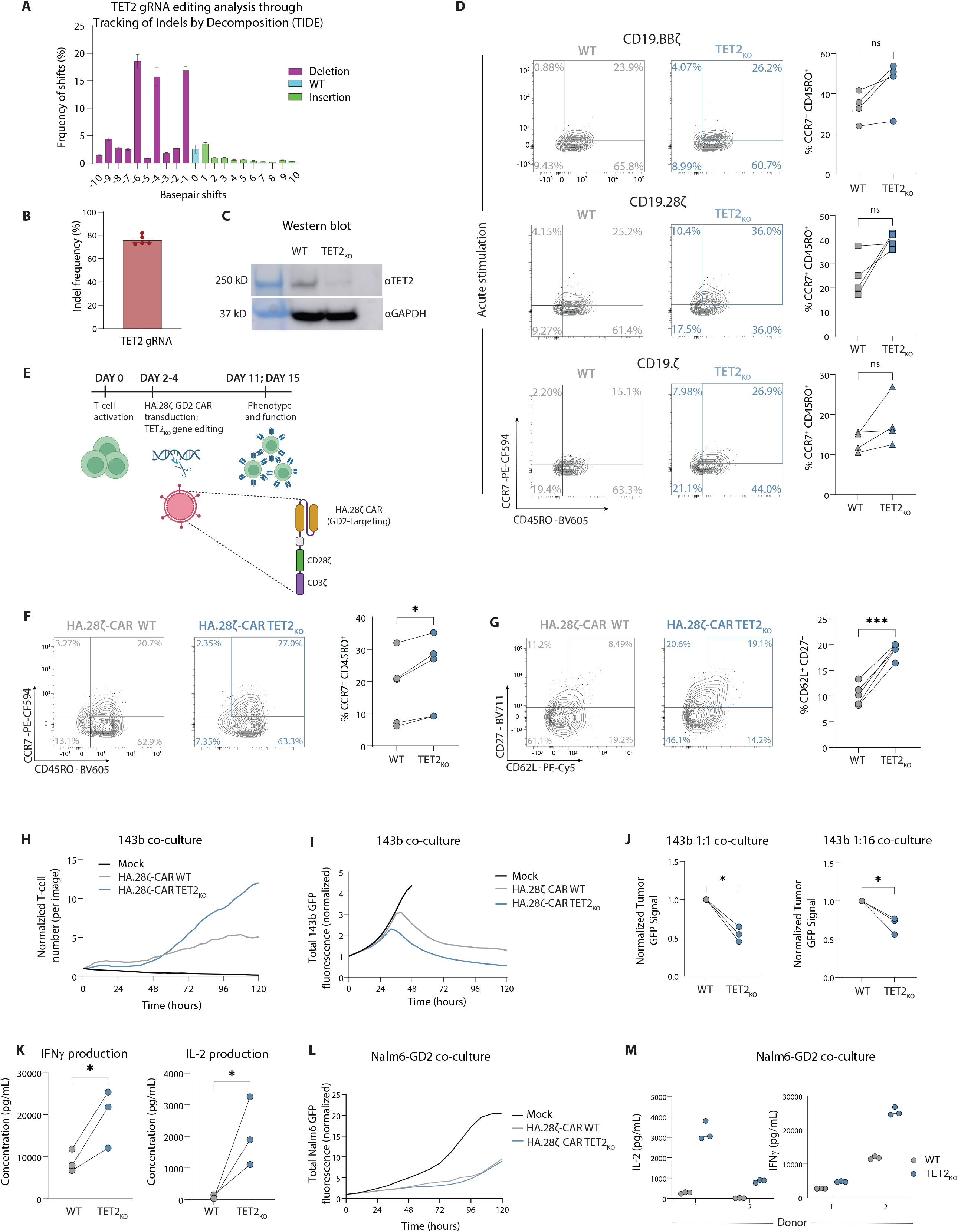
Validation of *TET2_KO_* and function of *TET2-*disrupted CAR T-cell during acute stimulation and in a tonic CAR signaling model. (**A-B**) Percent of sequences with indicated number of base pair shifts after CRISPR knockout by Tracking of Indels by Decomposition (TIDE) analysis (**A**). Average indel frequency in bulk CRISPR-edited populations via TIDE (**B**), n = 5. (**C**) Western blot showing loss of TET2 protein in *TET2_KO_* CAR T-cells. (**D**) Example plots and data of CD19.CD3ζ, CD19.BBζ, and CD19.CD28ζ CAR T-cells ± *TET2_KO_* showing distribution of CCR7^+^ CD45RO^+^ central memory-associated markers in CD8^+^ populations after 1 stimulation (acute stimulation), n = 4. (**E**) Schematic of GD2-targeting HA.28ζ CAR T-cell manufacturing and *TET2* gene editing. (**F-G**) Example plots and data of CD8^+^ CCR7^+^CD45RO^+^ (**F**) and CD27^+^CD62L^+^ (**G**) HA.28ζ CAR T-cell subsets ± *TET2_KO_* at day 11, n = 5. (**H**) HA.28ζ CAR T-cell ± *TET2_KO_* growth in co-culture with 143b-GL osteosarcoma tumor cells. (**I**) Tumor GFP florescence intensity after HA.28ζ CAR T-cell ± *TET2_KO_* co-culture with 143b-GL osteosarcoma tumor cells, run in triplicate. (**J**) Normalized tumor GFP signal from 1:1 and 1:16 E:T co-culture with 143b-GL tumor cells, n = 3. (**K**) IFNγ and IL-2 production after HA.28ζ CAR T-cell ± *TET2_KO_* co-culture with 143b-GL tumor cells, n = 3. (**L**) Tumor GFP florescence intensity after HA.28ζ CAR T-cell ± *TET2_KO_* co-culture against Nalm6-GD2^+^ tumor cells, run in triplicate. (**M**) Cytokine production from HA.28ζ CAR T ± *TET2_KO_* against Nalm6-GD2^+^ tumor cells, n = 2, run in triplicate. Data shown as mean ± SEM (**A-B**) or individual values (**D, F, G, J, K**) from independent donors or as mean of technical replicates (**H, I, L**). ns p > 0.05; *p < 0.05; **p < 0.01; ***p < 0.001 by paired t-test.

**Supplemental Figure 3.**
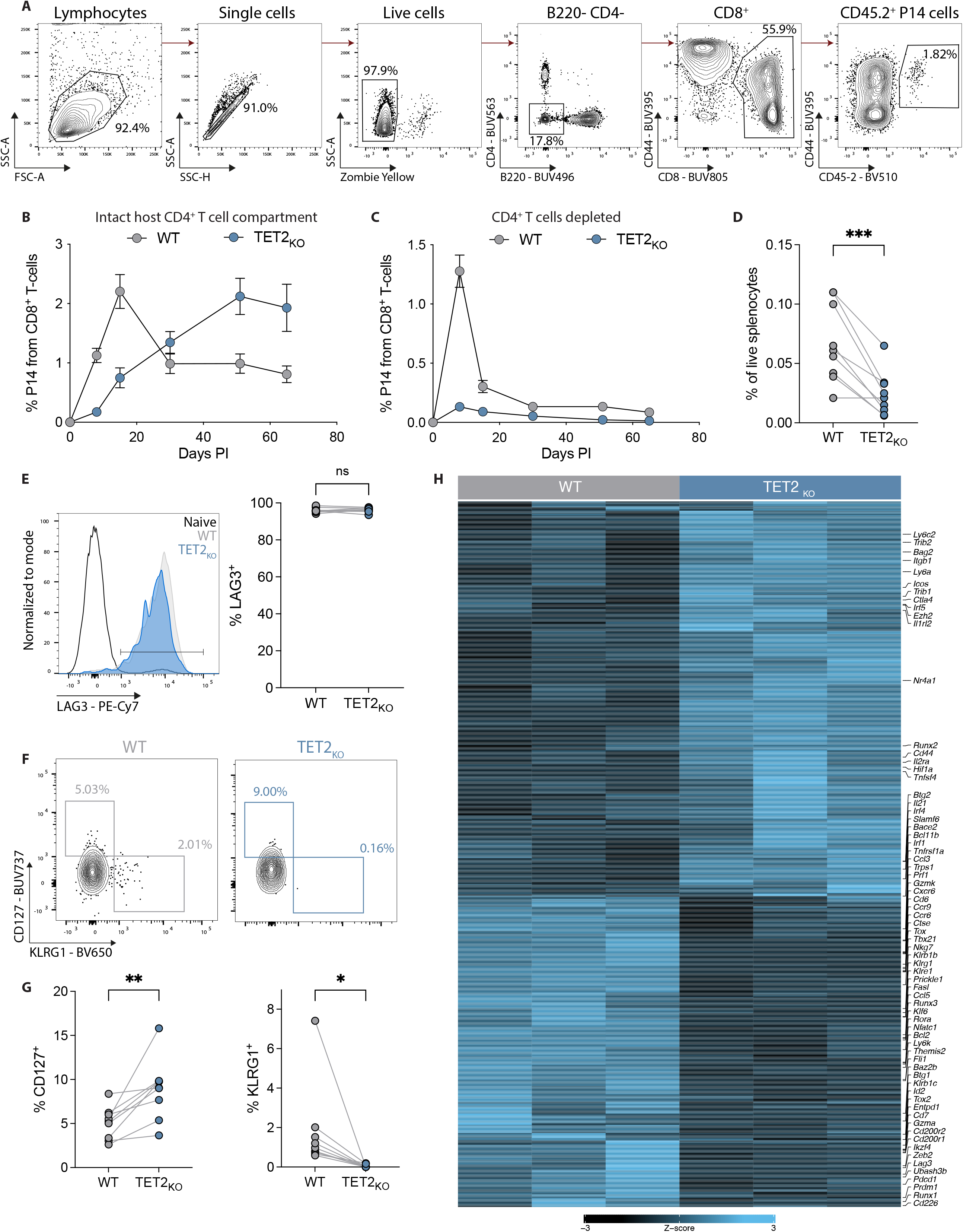
TET2 regulates differentiation of exhausted CD8^+^ T-cells. (**A**) Gating strategy for identification of co-transferred WT and *TET2_KO_* P14 cells. (**B**) Frequency of WT and *TET2_KO_* P14 cells in blood during chronic LCMV infection without CD4^+^ T-cell depletion. n = 15, mean ± SEM shown. Representative of 2 experiments. (**C**) Frequency of WT and *TET2_KO_* P14 cells in blood during chronic LCMV infection. Mice were treated at day −1 and day +1 with CD4-depleting antibody GK1.5, n = 8-10, mean ± SEM shown. Representative of at least 4 experiments. (**D**) Frequency of WT and *TET2_KO_* P14 cells in spleen. (**E**) Example plot and data showing LAG3 expression on WT compared to *TET2_KO_* P14 cells. (**F-G**) Example plots (**F**) and data (**G**) comparing KLRG1^+^ T_EFF_ and CD127^+^ T_MEM_ frequencies within WT and *TET2_KO_* P14 cells. (**H**) Heatmap comparing genes differentially expressed between WT and *TET2_KO_* P14 cells at day 15 p.i. with LCMV clone 13. (**D, E, F**) n = 9, spleen at day 30 p.i. with LCMV clone 13. Data for individual mice shown; representative of at least 4 experiments. ns p > 0.05; * p < 0.05; ** p < 0.01; *** p < 0.001; **** p < 0.0001 by paired t-test.

**Supplemental Figure 4.**
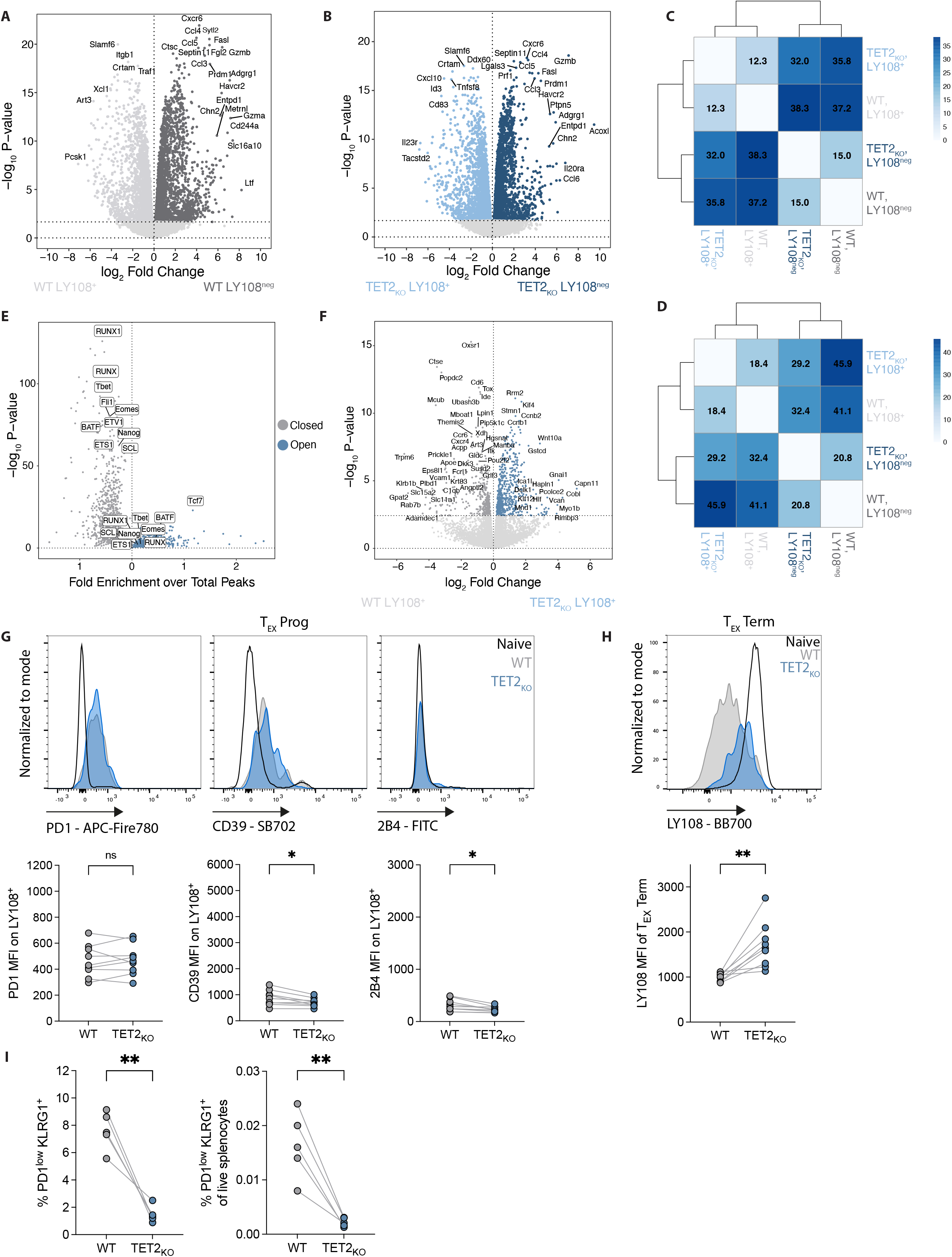
TET2 loss alters terminal T_EX_ differentiation. (**A-B**) Volcano plot highlighting DEGs between LY108^+^ and LY108^neg^ WT P14 cells (**A**) or LY108^+^ and LY108^neg^ *TET2_KO_* P14 cells (**B**). Dotted line at log_10_ P-value 0.05. (**C-D**) Distance analysis for RNA-seq (**C**) and ATAC-seq (**D**) data comparing T_EX_ subsets between WT and *TET2_KO_* P14 cells. (**E**) Volcano plot showing changes in transcription factor accessibility in WT LY108^neg^ cells versus *TET2_KO_* LY108^neg^ cells. x-axis represents fold enrichment over total peaks; y-axis represents -log_10_ P-value. (**F**) Volcano plot highlighting DEG in WT compared to *TET2_KO_* LY108^+^ T_EX_ cells. (**G**) Example plots and data comparing expression of PD1, CD39 and 2B4 on *TET2_KO_* LY108^+^ T_EX_ cells to WT LY108^+^ T_EX_ cells. (**H**) Example plots and data comparing expression of LY108 on *TET2_KO_* to WT terminally exhausted T_EX_. (**I**) Proportion and absolute frequency of PD1^low^ KLRG1^+^ T_EFF_-like WT and *TET2_KO_* P14 cells in spleen at day 6 p.i. with LCMV clone 13, n = 5. (**G-H**) n = 9, spleen at day 30 p.i. with LCMV clone 13. (**G, H, I**) Data for individual mice shown; representative of 2-3 independent experiments. * p < 0.05; ** p < 0.01 by paired t-test.

**Supplemental Figure 5.**
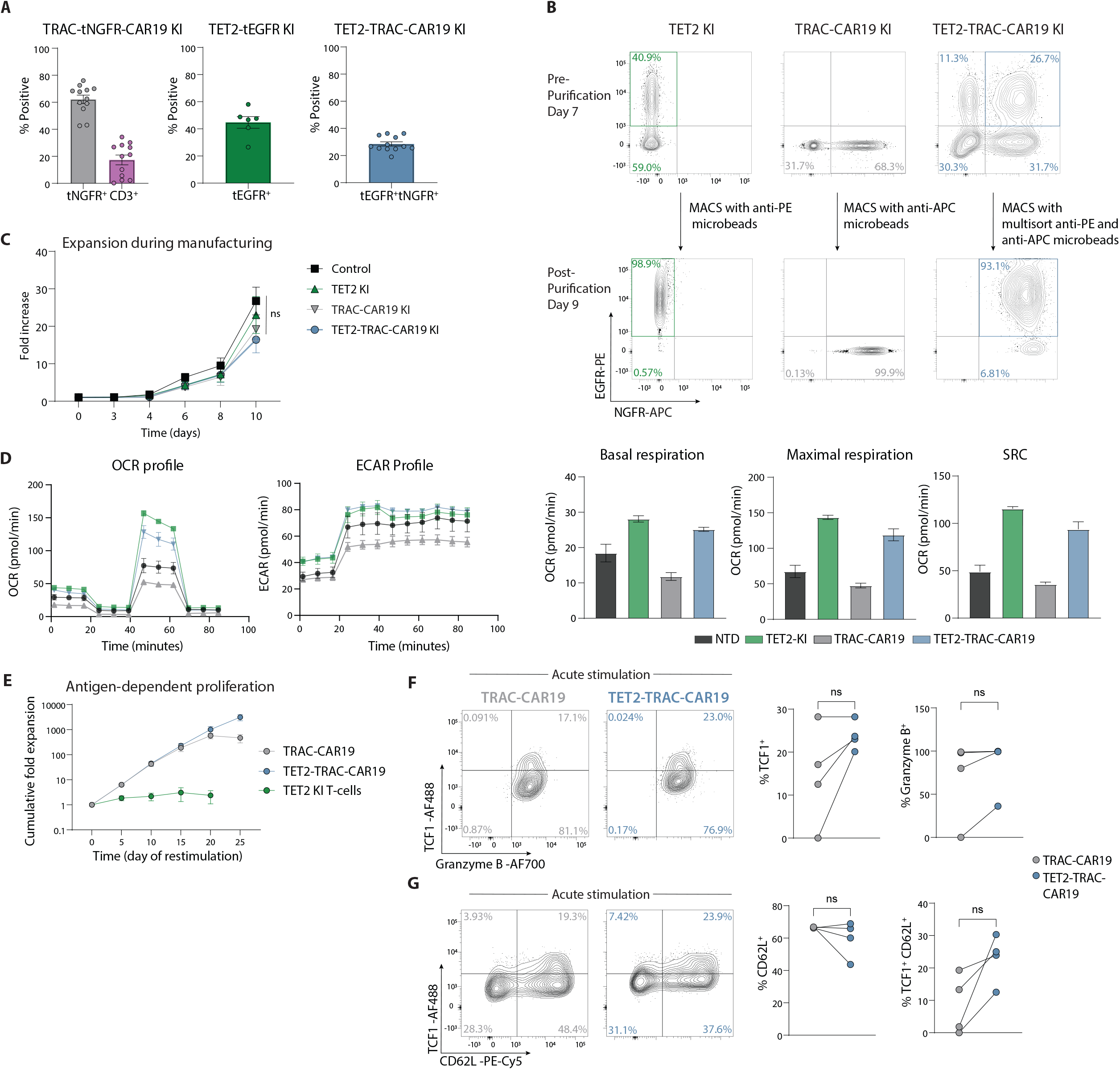
Engineering of TET2-TRAC-CAR19 dual knock-in T-cells. (**A**) Summary of TRAC-tNGFR knock-in (KI), CD3 knock-out (KO), TET2-tEGFR KI and TET2-TRAC-CAR19 KI editing efficiencies at day 7 during expansion, n = 6 – 12. (**B**) Example plots showing TET2-KI, TRAC-CAR19-KI and TET2-TRAC-CAR19 KI cells pre-(day 7) and post-purification (day 9). (**C**) T-cell expansion during manufacturing and CRISPR-AAV editing; mean ± SEM shown, n = 4, ns p > 0.05 by one-way ANOVA test with a post hoc Tukey’s multiple comparison test. (**D**) OCR and ECAR, basal respiration, maximal respiration and SRC profiles of NTD, TET2-KI, TRAC-CAR19 and TET2-TRAC-CAR19 T-cells at the end of manufacturing, n = 1, run in triplicate. (**E**) Antigen-dependent proliferation of TET2-TRAC-CAR T-cells compared to TET2-KI T-cells, n = 5. (**F**) Example plots and data of TCF1 and granzyme B expression on CD8^+^ CAR T-cells after 1 stimulation (acute stimulation), n = 4. (**G**) Example plots and data of CD8^+^ TCF1 and CD62L expression on CD8^+^ CAR T-cells after 1 stimulation (acute stimulation), n = 4. Data shown as mean ± SEM (**C**, **E**) or individual values (**A**, **F**, **G**) from independent donors or as mean ± SEM of technical replicates (**D**). (**F**, **G**) ns p > 0.05 by paired t-test.

**Supplemental Figure 6.**
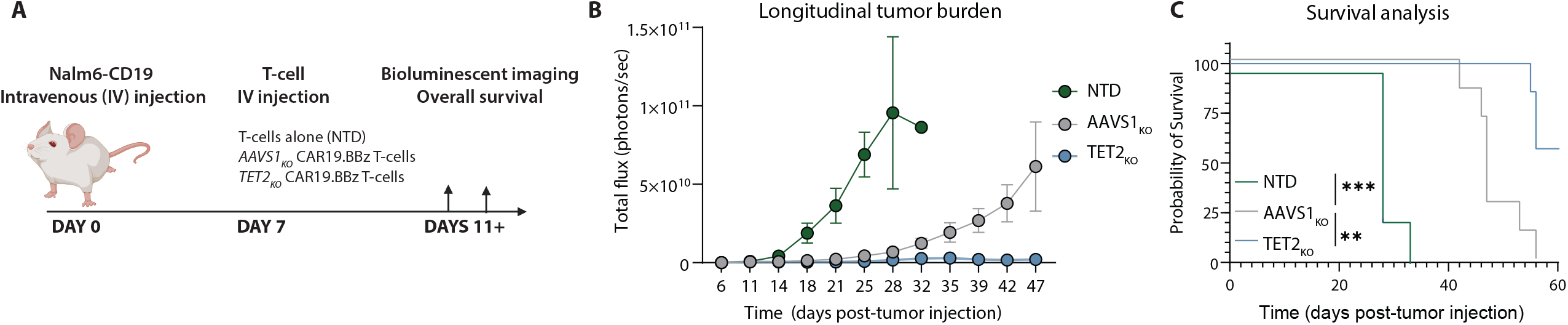
*In Vivo* tumor response of *TET2*_KO_ CAR T-cells. (**A**) Overview of *in vivo* experimental design, n = 4-7 mice per experimental group; data representative of one experiment. (**B**) Longitudinal analysis of *AAVS1*_KO_ and *TET2*_KO_ CAR T-cell group tumor burden compared to mice receiving non-transduced (NTD) T-cells. Data shown as mean ± SEM. (**C**) Overall group survival, ** p < 0.01; *** p < 0.001 by Mantel-Cox log-rank test for survival analysis.

**Table S1.** Recovered Integrations within 50KB of the *TET2* locus in CLL patient cohort.

| Patient | Cell Type | Time Point | Genomic Position | Relative Abundance |
| --- | --- | --- | --- | --- |
| p03712-12 | T-cells | Preinfusion | chr4-105264619 | 0.05 |
| p03712-16 | T-cells | Preinfusion | chr4-105225145 | 0.12 |
| p03712-29 | T-cells | Preinfusion | chr4+105291992 | 0.03 |
| p03712-31 | T-cells | Preinfusion | chr4+105183562 | 0.01 |
| p03712-40 | T-cells | Preinfusion | chr4-105163547 | 0.01 |
| p03712-40 | T-cells | Preinfusion | chr4+105182603 | 0.01 |
| p03712-43 | T-cells | Preinfusion | chr4+105190119 | 0.02 |
| p03712-48 | T-cells | Preinfusion | chr4+105223086 | 0.02 |
| p03712-55 | T-cells | Preinfusion | chr4+105221969 | 0.04 |
| p03712-57 | T-cells | Preinfusion | chr4-105187373 | 0.06 |
| p03712-60 | T-cells | Preinfusion | chr4-105223721 | 0.06 |
| p03712-60 | T-cells | Preinfusion | chr4+105265365 | 0.03 |
| p04409-06 | T-cells | Preinfusion | chr4-105224248 | 0.02 |
| p04409-07 | T-cells | Preinfusion | chr4+105267927 | 0.08 |
| p04409-09 | PBMC | Preinfusion | chr4+105253503 | 0.14 |
| p03712-03 | T-cells | First 15 Days | chr4+105160930 | 0.01 |
| p03712-03 | T-cells | First 15 Days | chr4+105200164 | 0.01 |
| p03712-03 | T-cells | First 15 Days | chr4-105298816 | 0.01 |
| p03712-18 | Whole Blood | First 15 Days | chr4-105248829 | 0.04 |
| p03712-18 | Whole Blood | First 15 Days | chr4+105153241 | 0.07 |
| p03712-18 | Whole Blood | First 15 Days | chr4+105251281 | 0.04 |
| p03712-18 | Whole Blood | First 15 Days | chr4+105260695 | 0.04 |
| p03712-18 | Whole Blood | First 15 Days | chr4-105224201 | 0.03 |
| p03712-29 | Whole Blood | First 15 Days | chr4+105228115 | 0.14 |
| p04409-09* | T-cells | First 15 Days | chr4+105190185 | 0.02 |
| p04409-09 | T-cells | First 15 Days | chr4+105257004 | 0.02 |
| p04409-22 | Whole Blood | First 15 Days | chr4-105152755 | 0.03 |
| p04409-22 | Whole Blood | First 15 Days | chr4-105164454 | 0.03 |
| p04409-22 | Whole Blood | First 15 Days | chr4+105263836 | 0.05 |
| p04409-09 | PBMC | Second 15 Days | chr4+105206047 | 0.23 |
| p04409-09* | Whole Blood | Second 15 Days | chr4+105190185 | 0.03 |
| p04409-09 | Bone Marrow | Second 15 Days | chr4-105230899 | 0.03 |
| p04409-22 | Bone Marrow | Second 15 Days | chr4+105164450 | 0.17 |
| p03712-47 | Bone Marrow | >30 Days | chr4-105155581 | 0.53 |

**Table S2.** Recovered Integrations within 50KB of the *TET2* locus in ALL patient cohort.

| Patient | Cell Type | Time Point | Genomic Position | Relative Abundance |
| --- | --- | --- | --- | --- |
| p959-101 | T-cells | Preinfusion | chr4-105122207 | 0.02 |
| p959-103 | T-cells | Preinfusion | chr4-105268362 | 0.04 |
| p959-103 | T-cells | Preinfusion | chr4+105197328 | 0.08 |
| p959-104 | T-cells | Preinfusion | chr4-105205781 | 0.05 |
| p959-104 | T-cells | Preinfusion | chr4+105165499 | 0.05 |
| p959-107 | T-cells | Preinfusion | chr4-105229630 | 0.01 |
| p959-107 | T-cells | Preinfusion | chr4+105151872 | 0.01 |
| p959-108 | T-cells | Preinfusion | chr4-105243661 | 0.06 |
| p959-111 | T-cells | Preinfusion | chr4+105219760 | 0.02 |
| p959-112 | T-cells | Preinfusion | chr4-105226123 | 0.02 |
| p959-112 | T-cells | Preinfusion | chr4+105151786 | 0.02 |
| p959-112 | T-cells | Preinfusion | chr4+105236100 | 0.04 |
| p959-113 | T-cells | Preinfusion | chr4-105158827 | 0.01 |
| p959-113 | T-cells | Preinfusion | chr4+105221551 | 0.01 |
| p959-113 | T-cells | Preinfusion | chr4+105271906 | 0.01 |
| p959-115 | T-cells | Preinfusion | chr4-105229150 | 0.02 |
| p959-117 | T-cells | Preinfusion | chr4+105173039 | 0.03 |
| p959-118 | T-cells | Preinfusion | chr4-105162564 | 0.01 |
| p959-118 | T-cells | Preinfusion | chr4-105204835 | 0.01 |
| p959-118 | T-cells | Preinfusion | chr4-105227631 | 0.01 |
| p959-118 | T-cells | Preinfusion | chr4-105241238 | 0.01 |
| p959-118 | T-cells | Preinfusion | chr4+105182278 | 0.01 |
| p959-118 | T-cells | Preinfusion | chr4+105224901 | 0.01 |
| p959-121 | T-cells | Preinfusion | chr4-105204911 | 0.04 |
| p959-123 | T-cells | Preinfusion | chr4-105239813 | 0.01 |
| p959-123 | T-cells | Preinfusion | chr4+105171995 | 0.04 |
| p959-123 | T-cells | Preinfusion | chr4+105224760 | 0.03 |
| p959-123 | T-cells | Preinfusion | chr4+105243003 | 0.01 |
| p959-125 | T-cells | Preinfusion | chr4-105231290 | 0.02 |
| p959-131 | T-cells | Preinfusion | chr4-105225782 | 0.03 |
| p959-131 | T-cells | Preinfusion | chr4+105187755 | 0.03 |
| p959-132 | T-cells | Preinfusion | chr4+105250821 | 0.07 |
| p959-132 | T-cells | Preinfusion | chr4+105260601 | 0.07 |
| p959-133 | T-cells | Preinfusion | chr4+105256735 | 0.02 |
| p959-133 | T-cells | Preinfusion | chr4-105260182 | 0.02 |
| p959-136 | T-cells | Preinfusion | chr4-105186871 | 0.03 |
| p959-136 | T-cells | Preinfusion | chr4+105157216 | 0.03 |
| p959-136 | T-cells | Preinfusion | chr4+105203882 | 0.03 |
| p959-141 | T-cells | Preinfusion | chr4+105228898 | 0.03 |
| p959-141 | T-cells | Preinfusion | chr4+105294015 | 0.03 |
| p959-144 | T-cells | Preinfusion | chr4+105175287 | 0.02 |
| p959-144 | T-cells | Preinfusion | chr4+105202132 | 0.02 |
| p959-146 | T-cells | Preinfusion | chr4+105154691 | 0.02 |
| p959-157 | T-cells | Preinfusion | chr4+105243230 | 0.03 |
| p959-158 | T-cells | Preinfusion | chr4-105160134 | 0.01 |
| p959-158 | T-cells | Preinfusion | chr4+105227059 | 0.01 |
| p959-169 | T-cells | Preinfusion | chr4+105225389 | 0.04 |
| p959-172 | T-cells | Preinfusion | chr4+105273807 | 0.05 |
| p959-174 | T-cells | Preinfusion | chr4-105204656 | 0.08 |
| p959-158 | T-cells | Preinfusion | chr4-105191937 | 0.01 |
| p959-158 | T-cells | Preinfusion | chr4-105257743 | 0.01 |
| p959-158 | T-cells | Preinfusion | chr4-105271578 | 0.01 |
| p959-158 | T-cells | Preinfusion | chr4+105284604 | 0.01 |
| p959-101 | Whole Blood | First 15 Days | chr4+105206388 | 0.05 |
| p959-105 | Whole Blood | First 15 Days | chr4+105231121 | 0.06 |
| p959-113 | PBMC | First 15 Days | chr4+105230717 | 0.02 |
| p959-121 | Whole Blood | First 15 Days | chr4+105223180 | 0.08 |
| p959-125 | Whole Blood | First 15 Days | chr4+105229646 | 0.02 |
| p959-125 | Whole Blood | First 15 Days | chr4+105258957 | 0.02 |
| p959-139 | Whole Blood | First 15 Days | chr4-105275946 | 0.07 |
| p959-139 | Whole Blood | First 15 Days | chr4+105142570 | 0.03 |
| p959-140 | Whole Blood | First 15 Days | chr4-105269087 | 0.03 |
| p959-141 | Whole Blood | First 15 Days | chr4-105201535 | 0.07 |
| p959-146 | Whole Blood | First 15 Days | chr4-105185661 | 0.04 |
| p959-160 | Whole Blood | First 15 Days | chr4+105217644 | 0.02 |
| p959-160 | Whole Blood | First 15 Days | chr4+105262568 | 0.02 |
| p04409-29 | PBMC | Second 15 Days | chr4+105269578 | 0.08 |
| p959-143 | Whole Blood | Second 15 Days | chr4-105259704 | 0.05 |
| p959-153 | Bone Marrow | Second 15 Days | chr4-105280290 | 0.10 |
| p959-160 | Whole Blood | Second 15 Days | chr4-105208249 | 0.03 |
| p959-160 | Whole Blood | Second 15 Days | chr4+105227846 | 0.03 |
| p959-160 | Whole Blood | Second 15 Days | chr4+105262098 | 0.03 |
| p959-160 | Bone Marrow | Second 15 Days | chr4-105247460 | 0.05 |
| p959-100 | Bone Marrow | >30 Days | chr4-105198703 | 0.33 |
| p959-160 | Whole Blood | >30 Days | chr4-105238384 | 0.46 |

**Table S6.**
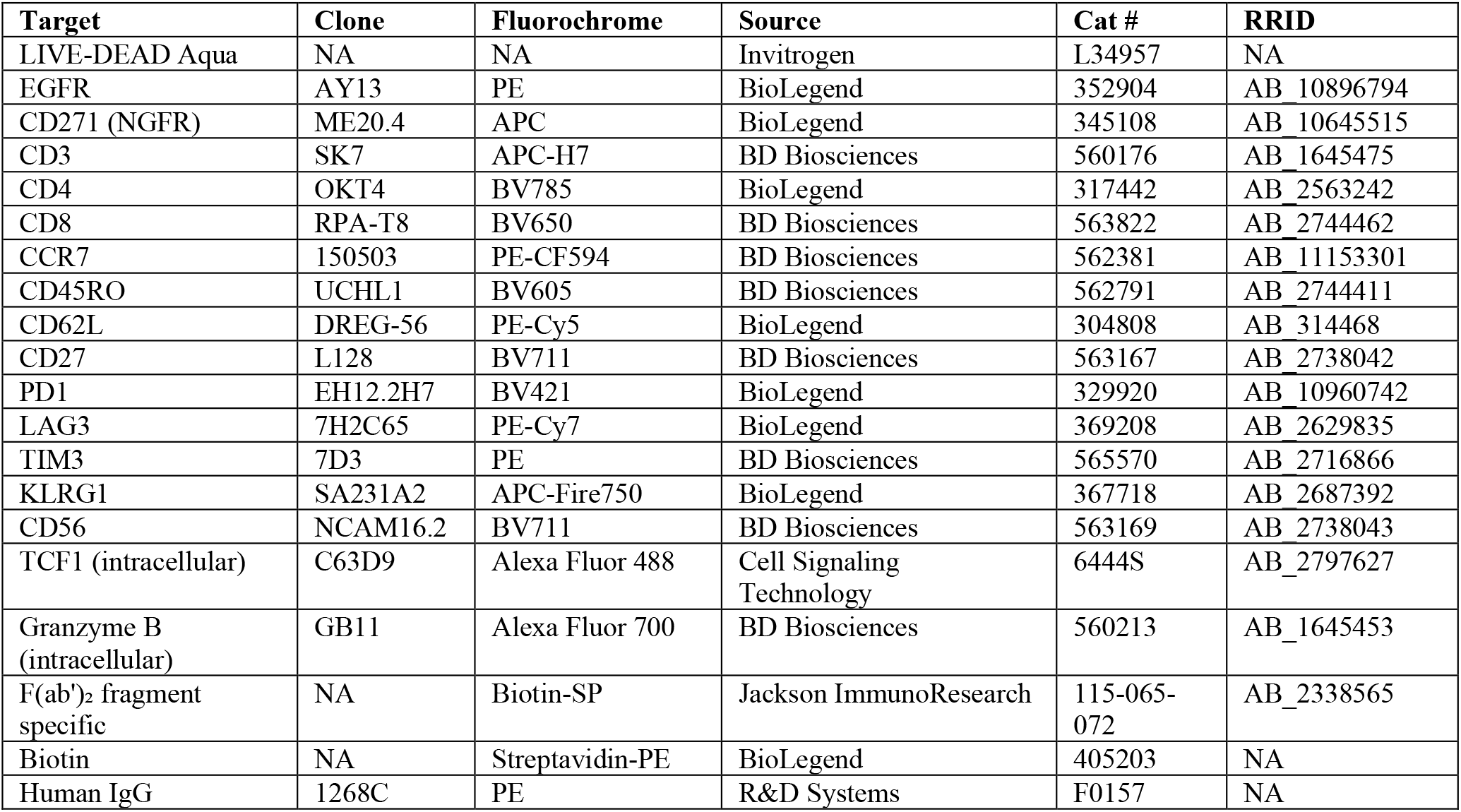
Antibodies for CAR T-cell studies.

| Target | Clone | Fluorochrome | Source | Cat # | RRID |
| --- | --- | --- | --- | --- | --- |
| LIVE-DEAD Aqua | NA | NA | Invitrogen | L34957 | NA |
| EGFR | AY13 | PE | BioLegend | 352904 | AB_10896794 |
| CD271 (NGFR) | ME20.4 | APC | BioLegend | 345108 | AB_10645515 |
| CD3 | SK7 | APC-H7 | BD Biosciences | 560176 | AB_1645475 |
| CD4 | OKT4 | BV785 | BioLegend | 317442 | AB_2563242 |
| CD8 | RPA-T8 | BV650 | BD Biosciences | 563822 | AB_2744462 |
| CCR7 | 150503 | PE-CF594 | BD Biosciences | 562381 | AB_11153301 |
| CD45RO | UCHL1 | BV605 | BD Biosciences | 562791 | AB_2744411 |
| CD62L | DREG-56 | PE-Cy5 | BioLegend | 304808 | AB_314468 |
| CD27 | L128 | BV711 | BD Biosciences | 563167 | AB_2738042 |
| PD1 | EH12.2H7 | BV421 | BioLegend | 329920 | AB_10960742 |
| LAG3 | 7H2C65 | PE-Cy7 | BioLegend | 369208 | AB_2629835 |
| TIM3 | 7D3 | PE | BD Biosciences | 565570 | AB_2716866 |
| KLRG1 | SA231A2 | APC-Fire750 | BioLegend | 367718 | AB_2687392 |
| CD56 | NCAM16.2 | BV711 | BD Biosciences | 563169 | AB_2738043 |
| TCF1 (intracellular) | C63D9 | Alexa Fluor 488 | Cell Signaling Technology | 6444S | AB_2797627 |
| Granzyme B (intracellular) | GB11 | Alexa Fluor 700 | BD Biosciences | 560213 | AB_1645453 |
| F(ab')<sub>2</sub> fragment specific | NA | Biotin-SP | Jackson ImmunoResearch | 115-065-072 | AB_2338565 |
| Biotin | NA | Streptavidin-PE | BioLegend | 405203 | NA |
| Human IgG | 1268C | PE | R&D Systems | F0157 | NA |

**Table S7.** Antibodies for LCMV studies.

| Target | Clone | Fluorochrome | Source | Cat # | RRID |
| --- | --- | --- | --- | --- | --- |
| LiveDead | NA | AquaVivid | ThermoFisher | L34957 | NA |
| LiveDead | NA | Zombie Yellow | BioLegend | 423104 | NA |
| LiveDead | NA | Zombie NIR | BioLegend | 423106 | NA |
| B220 | RA3-6B2 | APC-eF780 | ThermoFisher | 47-0452-82 | AB_1518810 |
| B220 | RA3-6B2 | BUV496 | BD Biosciences | 612950 | AB_2870227 |
| CD4 | RM4-5 | APC-eF780 | ThermoFisher | 47-0042-82 | AB_1272183 |
| CD4 | GK1.5 | BUV563 | BD Biosciences | 612923 | AB_2870208 |
| CD8a | 53-6.7 | BUV805 | BD Biosciences | 612898 | AB_2870186 |
| CD8a | 53-6.7 | BV650 | BD Biosciences | 563234 | AB_2738084 |
| CD8a | 53-6.7 | APC | Invitrogen | 17-0081-83 | AB_469336 |
| CD39 | 24DMS1 | SB702 | Invitrogen | 67-0391-82 | AB_2717143 |
| CD44 | IM7 | BUV396 | BD Biosciences | 740215 | AB_2739963 |
| CD44 | IM7 | BV711 | BioLegend | 103057 | AB_2564214 |
| CD44 | IM7 | BV785 | BioLegend | 103059 | AB_2571953 |
| CD44 | IM7 | APC | BD Biosciences | 559250 | AB_398661 |
| CD45.1 | A20 | Pacific Blue | BioLegend | 110722 | AB_492866 |
| CD45.1 | A20 | PE-Cy5 | ThermoFisher | 15-0453-81 | AB_468758 |
| CD45.1 | A20 | AF700 | BioLegend | 110724 | AB_493733 |
| CD45.2 | 104 | BV570 | BioLegend | 109833 | AB_10900987 |
| CD45.2 | 104 | BV785 | BioLegend | 109839 | AB_2562604 |
| CD45.2 | 104 | AF700 | BioLegend | 109822 | AB_493731 |
| CD62L | MEL-14 | BV605 | BD Biosciences | 563252 | AB_2738098 |
| CD62L | MEL-14 | PE-CF594 | BD Biosciences | 562404 | AB_11154046 |
| CD69 | H1.2F3 | BV480 | BD Biosciences | 746813 | AB_2744067 |
| CD127 | SB/199 | BUV737 | BD Biosciences | 612841 | AB_2870163 |
| CD127 | SB/199 | PE-CF594 | BD Biosciences | 562419 | AB_11153131 |
| CD244 (2B4) | eBio244F4 | FITC | eBioscience | 11-2441-85 | AB_657877 |
| CD244 (2B4) | 2B4 | FITC | BD Biosciences | 553305 | AB_394769 |
| CX3CR1 | SA011F11 | BV605 | BioLegend | 149027 | AB_2565937 |
| CX3CR1 | SA011F11 | BV650 | BioLegend | 149033 | AB_2565999 |
| CXCR3 | CXCR3-173 | Biotin | BioLegend | 126503 | AB_1027658 |
| GFP | Polyclonal | AF488 | ThermoFisher | A-21311 | AB_221477 |
| gp33 | NA | PE | In house | NA | NA |
| Granzyme B | GB11 | BV421 | BD Biosciences | 563389 | AB_2738175 |
| IFNγ | XMG1.2 | PerCP-Cy5.5 | BioLegend | 505822 | AB_961361 |
| KI67 | B56 | A700 | BD Biosciences | 561277 | AB_10611571 |
| KLRG1 | 2F1 | FITC | eBioscience | 11-5893-80 | AB_1311268 |
| KLRG1 | 2F1 | PerCP-eF710 | Invitrogen | 46-5893-82 | AB_10670282 |
| KLRG1 | 2F1 | BV650 | BD Biosciences | 740553 | AB_2740254 |
| LAG3 | C9B7W | PE-Cy7 | BioLegend | 125226 | AB_2715764 |
| LAG3 | C9B7W | AF647 | BioRad | MCA2386A647T | AB_2133342 |
| LY108 | 13G3 | BV421 | BD Biosciences | 740090 | AB_2739850 |
| LY108 | 13G3 | BB700 | BD Biosciences | 742272 | AB_2871448 |
| PD-1 | RMPI-30 | PE-Cy7 | BioLegend | 109110 | AB_572017 |
| PD-1 | 29F.1A12 | APC-Fire780 | BioLegend | 135239 | AB_2629767 |
| TCF-1 | S33-966 | PE | BD Biosciences | 564217 | AB_2687845 |
| TIGIT | 1G9 | BV421 | BD Biosciences | 565270 | AB_2688007 |
| TIGIT | 1G9 | PE-Dazzle594 | BioLegend | 142110 | AB_2566573 |
| TIM3 | RMT3-23 | BV605 | BioLegend | 119721 | AB_2616907 |
| TIM3 | RMT3-23 | BV785 | BioLegend | 119725 | AB_2716066 |
| TNF-alpha | MP6-XT22 | Pacific Blue | BioLegend | 506318 | AB_893639 |
| TOX | REA473 | APC | Miltenyi | 130-118-335 | AB_2751485 |

## REFERENCES AND NOTES

1. C. U. Blank, W. N. Haining, W. Held, P. G. Hogan, A. Kallies, E. Lugli, R. C. Lynn, M. Philip, A. Rao, N. P. Restifo, A. Schietinger, T. N. Schumacher, P. L. Schwartzberg, A. H. Sharpe, D. E. Speiser, E. J. Wherry, B. A. Youngblood, D. Zehn, Defining ‘T cell exhaustion’. Nat Rev Immunol 19, 665–674 (2019).

2. S. M. Albelda, CAR T cell therapy for patients with solid tumours: key lessons to learn and unlearn. Nat Rev Clin Oncol 21, 47–66 (2024).

3. R. C. Lynn, E. W. Weber, E. Sotillo, D. Gennert, P. Xu, Z. Good, H. Anbunathan, J. Lattin, R. Jones, V. Tieu, S. Nagaraja, J. Granja, C. F. A. de Bourcy, R. Majzner, A. T. Satpathy, S. R. Quake, M. Monje, H. Y. Chang, C. L. Mackall, c-Jun overexpression in CAR T cells induces exhaustion resistance. Nature 576, 293–300 (2019).

4. A. H. Long, W. M. Haso, J. F. Shern, K. M. Wanhainen, M. Murgai, M. Ingaramo, J. P. Smith, A. J. Walker, M. E. Kohler, V. R. Venkateshwara, R. N. Kaplan, G. H. Patterson, T. J. Fry, R. J. Orentas, C. L. Mackall, 4-1BB costimulation ameliorates T cell exhaustion induced by tonic signaling of chimeric antigen receptors. Nat Med 21, 581–590 (2015).

5. J. Chen, I. F. Lopez-Moyado, H. Seo, C. J. Lio, L. J. Hempleman, T. Sekiya, A. Yoshimura, J. P. Scott-Browne, A. Rao, NR4A transcription factors limit CAR T cell function in solid tumours. Nature 567, 530–534 (2019).

6. I. Y. Jung, V. Narayan, S. McDonald, A. J. Rech, R. Bartoszek, G. Hong, M. M. Davis, J. Xu, A. C. Boesteanu, J. S. Barber-Rotenberg, G. Plesa, S. F. Lacey, J. K. Jadlowsky, D. L. Siegel, D. M. Hammill, P. F. Cho-Park, S. L. Berger, N. B. Haas, J. A. Fraietta, BLIMP1 and NR4A3 transcription factors reciprocally regulate antitumor CAR T cell stemness and exhaustion. Sci Transl Med 14, eabn7336 (2022).

7. P. S. Adusumilli, M. G. Zauderer, I. Riviere, S. B. Solomon, V. W. Rusch, R. E. O’Cearbhaill, A. Zhu, W. Cheema, N. K. Chintala, E. Halton, J. Pineda, R. Perez-Johnston, K. S. Tan, B. Daly, J. A. Araujo Filho, D. Ngai, E. McGee, A. Vincent, C. Diamonte, J. L. Sauter, S. Modi, D. Sikder, B. Senechal, X. Wang, W. D. Travis, M. Gonen, C. M. Rudin, R. J. Brentjens, D. R. Jones, M. Sadelain, A Phase I Trial of Regional Mesothelin-Targeted CAR T-cell Therapy in Patients with Malignant Pleural Disease, in Combination with the Anti-PD-1 Agent Pembrolizumab. Cancer Discov 11, 2748–2763 (2021).

8. K. E. Pauken, M. A. Sammons, P. M. Odorizzi, S. Manne, J. Godec, O. Khan, A. M. Drake, Z. Chen, D. R. Sen, M. Kurachi, R. A. Barnitz, C. Bartman, B. Bengsch, A. C. Huang, J. M. Schenkel, G. Vahedi, W. N. Haining, S. L. Berger, E. J. Wherry, Epigenetic stability of exhausted T cells limits durability of reinvigoration by PD-1 blockade. Science 354, 1160–1165 (2016).

9. B. Prinzing, C. C. Zebley, C. T. Petersen, Y. Fan, A. A. Anido, Z. Yi, P. Nguyen, H. Houke, M. Bell, D. Haydar, C. Brown, S. K. Boi, S. Alli, J. C. Crawford, J. M. Riberdy, J. J. Park, S. Zhou, M. P. Velasquez, C. DeRenzo, C. R. Lazzarotto, S. Q. Tsai, P. Vogel, S. M. Pruett-Miller, D. M. Langfitt, S. Gottschalk, B. Youngblood, G. Krenciute, Deleting DNMT3A in CAR T cells prevents exhaustion and enhances antitumor activity. Sci Transl Med 13, eabh0272 (2021).

10. J. A. Fraietta, C. L. Nobles, M. A. Sammons, S. Lundh, S. A. Carty, T. J. Reich, A. P. Cogdill, J. J. D. Morrissette, J. E. Denizio, S. Reddy, Y. Hwang, M. Gohil, I. Kulikovskaya, F. Nazimuddin, M. Gupta, F. Chen, J. K. Everett, K. A. Alexander, E. Lin-Shiao, …, J. J. Melenhorst, Disruption of TET2 promotes the therapeutic efficacy of CD19-targeted T cells. Nature 558, 307–312 (2018).

11. K. D. Rasmussen, K. Helin, Role of TET enzymes in DNA methylation, development, and cancer. Genes & Development 30, 733–750 (2016).

12. L. Tan, Y. G. Shi, Tet family proteins and 5-hydroxymethylcytosine in development and disease. Development 139, 1895–1902 (2012).

13. N. Jain, Z. Zhao, J. Feucht, R. Koche, A. Iyer, A. Dobrin, J. Mansilla-Soto, J. Yang, Y. Zhan, M. Lopez, G. Gunset, M. Sadelain, TET2 guards against unchecked BATF3-induced CAR T cell expansion. Nature 615, 315–322 (2023).

14. S. A. Carty, M. Gohil, L. B. Banks, R. M. Cotton, M. E. Johnson, E. Stelekati, A. D. Wells, E. J. Wherry, G. A. Koretzky, M. S. Jordan, The Loss of TET2 Promotes CD8(+) T Cell Memory Differentiation. J Immunol 200, 82–91 (2018).

15. M. Irving, E. Lanitis, D. Migliorini, Z. Ivics, S. Guedan, Choosing the Right Tool for Genetic Engineering: Clinical Lessons from Chimeric Antigen Receptor-T Cells. Human Gene Therapy 32, 1044–1058 (2021).

16. T. K. MacLachlan, in Nonclinical Development of Novel Biologics, Biosimilars, Vaccines and Specialty Biologics, L. M. Plitnick, D. J. Herzyk, Eds. (Academic Press, San Diego, 2013), pp. 259–285.

17. G. J. van der Windt, D. O’Sullivan, B. Everts, S. C. Huang, M. D. Buck, J. D. Curtis, C. H. Chang, A. M. Smith, T. Ai, B. Faubert, R. G. Jones, E. J. Pearce, E. L. Pearce, CD8 memory T cells have a bioenergetic advantage that underlies their rapid recall ability. Proc Natl Acad Sci U S A 110, 14336–14341 (2013).

18. B. Bengsch, A. L. Johnson, M. Kurachi, P. M. Odorizzi, K. E. Pauken, J. Attanasio, E. Stelekati, L. M. McLane, M. A. Paley, G. M. Delgoffe, E. J. Wherry, Bioenergetic Insufficiencies Due to Metabolic Alterations Regulated by the Inhibitory Receptor PD-1 Are an Early Driver of CD8(+) T Cell Exhaustion. Immunity 45, 358–373 (2016).

19. W. Kong, A. Dimitri, W. Wang, I. Y. Jung, C. J. Ott, M. Fasolino, Y. Wang, I. Kulikovskaya, M. Gupta, T. Yoder, J. E. DeNizio, J. K. Everett, E. F. Williams, J. Xu, J. Scholler, T. J. Reich, V. G. Bhoj, K. M. Haines, M. V. Maus, J. J. Melenhorst, R. M. Young, J. K. Jadlowsky, K. T. Marcucci, J. E. Bradner, B. L. Levine, D. L. Porter, F. D. Bushman, R. M. Kohli, C. H. June, M. M. Davis, S. F. Lacey, G. Vahedi, J. A. Fraietta, BET bromodomain protein inhibition reverses chimeric antigen receptor extinction and reinvigorates exhausted T cells in chronic lymphocytic leukemia. J Clin Invest 131, (2021).

20. G. Escobar, D. Mangani, A. C. Anderson, T cell factor 1: A master regulator of the T cell response in disease. Sci Immunol 5, (2020).

21. J. E. Wu, S. Manne, S. F. Ngiow, A. E. Baxter, H. Huang, E. Freilich, M. L. Clark, J. H. Lee, Z. Chen, O. Khan, R. P. Staupe, Y. J. Huang, J. Shi, J. R. Giles, E. J. Wherry, In vitro modeling of CD8(+) T cell exhaustion enables CRISPR screening to reveal a role for BHLHE40. Sci Immunol 8, eade3369 (2023).

22. E. J. Wherry, R. Ahmed, Memory CD8 T-cell differentiation during viral infection. J Virol 78, 5535–5545 (2004).

23. A. Gallimore, A. Glithero, A. Godkin, A. C. Tissot, A. Pluckthun, T. Elliott, H. Hengartner, R. Zinkernagel, Induction and exhaustion of lymphocytic choriomeningitis virus-specific cytotoxic T lymphocytes visualized using soluble tetrameric major histocompatibility complex class I-peptide complexes. J Exp Med 187, 1383–1393 (1998).

24. A. J. Zajac, J. N. Blattman, K. Murali-Krishna, D. J. Sourdive, M. Suresh, J. D. Altman, R. Ahmed, Viral immune evasion due to persistence of activated T cells without effector function. J Exp Med 188, 2205–2213 (1998).

25. D. Moskophidis, F. Lechner, H. Pircher, R. M. Zinkernagel, Virus persistence in acutely infected immunocompetent mice by exhaustion of antiviral cytotoxic effector T cells. Nature 362, 758–761 (1993).

26. J. M. Angelosanto, S. D. Blackburn, A. Crawford, E. J. Wherry, Progressive loss of memory T cell potential and commitment to exhaustion during chronic viral infection. J Virol 86, 8161–8170 (2012).

27. E. J. Wherry, V. Teichgraber, T. C. Becker, D. Masopust, S. M. Kaech, R. Antia, U. H. von Andrian, R. Ahmed, Lineage relationship and protective immunity of memory CD8 T cell subsets. Nat Immunol 4, 225–234 (2003).

28. S. M. Kaech, J. T. Tan, E. J. Wherry, B. T. Konieczny, C. D. Surh, R. Ahmed, Selective expression of the interleukin 7 receptor identifies effector CD8 T cells that give rise to long-lived memory cells. Nat Immunol 4, 1191–1198 (2003).

29. S. D. Blackburn, H. Shin, G. J. Freeman, E. J. Wherry, Selective expansion of a subset of exhausted CD8 T cells by alphaPD-L1 blockade. Proc Natl Acad Sci U S A 105, 15016–15021 (2008).

30. B. C. Miller, D. R. Sen, R. Al Abosy, K. Bi, Y. V. Virkud, M. W. LaFleur, K. B. Yates, A. Lako, K. Felt, G. S. Naik, M. Manos, E. Gjini, J. R. Kuchroo, J. J. Ishizuka, J. L. Collier, G. K. Griffin, S. Maleri, D. E. Comstock, S. A. Weiss, F. D. Brown, A. Panda, M. D. Zimmer, R. T. Manguso, F. S. Hodi, S. J. Rodig, A. H. Sharpe, W. N. Haining, Subsets of exhausted CD8(+) T cells differentially mediate tumor control and respond to checkpoint blockade. Nat Immunol 20, 326–336 (2019).

31. M. A. Paley, D. C. Kroy, P. M. Odorizzi, J. B. Johnnidis, D. V. Dolfi, B. E. Barnett, E. K. Bikoff, E. J. Robertson, G. M. Lauer, S. L. Reiner, E. J. Wherry, Progenitor and terminal subsets of CD8+ T cells cooperate to contain chronic viral infection. Science 338, 1220–1225 (2012).

32. S. J. Im, M. Hashimoto, M. Y. Gerner, J. Lee, H. T. Kissick, M. C. Burger, Q. Shan, J. S. Hale, J. Lee, T. H. Nasti, A. H. Sharpe, G. J. Freeman, R. N. Germain, H. I. Nakaya, H. H. Xue, R. Ahmed, Defining CD8+ T cells that provide the proliferative burst after PD-1 therapy. Nature 537, 417–421 (2016).

33. D. T. Utzschneider, M. Charmoy, V. Chennupati, L. Pousse, D. P. Ferreira, S. Calderon-Copete, M. Danilo, F. Alfei, M. Hofmann, D. Wieland, S. Pradervand, R. Thimme, D. Zehn, W. Held, T Cell Factor 1-Expressing Memory-like CD8(+) T Cells Sustain the Immune Response to Chronic Viral Infections. Immunity 45, 415–427 (2016).

34. J. C. Beltra, S. Manne, M. S. Abdel-Hakeem, M. Kurachi, J. R. Giles, Z. Chen, V. Casella, S. F. Ngiow, O. Khan, Y. J. Huang, P. Yan, K. Nzingha, W. Xu, R. K. Amaravadi, X. Xu, G. C. Karakousis, T. C. Mitchell, L. M. Schuchter, A. C. Huang, E. J. Wherry, Developmental Relationships of Four Exhausted CD8(+) T Cell Subsets Reveals Underlying Transcriptional and Epigenetic Landscape Control Mechanisms. Immunity 52, 825–841 e828 (2020).

35. I. Siddiqui, K. Schaeuble, V. Chennupati, S. A. Fuertes Marraco, S. Calderon-Copete, D. Pais Ferreira, S. J. Carmona, L. Scarpellino, D. Gfeller, S. Pradervand, S. A. Luther, D. E. Speiser, W. Held, Intratumoral Tcf1(+)PD-1(+)CD8(+) T Cells with Stem-like Properties Promote Tumor Control in Response to Vaccination and Checkpoint Blockade Immunotherapy. Immunity 50, 195–211 e110 (2019).

36. O. Khan, J. R. Giles, S. McDonald, S. Manne, S. F. Ngiow, K. P. Patel, M. T. Werner, A. C. Huang, K. A. Alexander, J. E. Wu, J. Attanasio, P. Yan, S. M. George, B. Bengsch, R. P. Staupe, G. Donahue, W. Xu, R. K. Amaravadi, X. Xu, G. C. Karakousis, T. C. Mitchell, L. M. Schuchter, J. Kaye, S. L. Berger, E. J. Wherry, TOX transcriptionally and epigenetically programs CD8(+) T cell exhaustion. Nature 571, 211–218 (2019).

37. H. Seo, J. Chen, E. Gonzalez-Avalos, D. Samaniego-Castruita, A. Das, Y. H. Wang, I. F. Lopez-Moyado, R. O. Georges, W. Zhang, A. Onodera, C. J. Wu, L. F. Lu, P. G. Hogan, A. Bhandoola, A. Rao, TOX and TOX2 transcription factors cooperate with NR4A transcription factors to impose CD8(+) T cell exhaustion. Proc Natl Acad Sci U S A 116, 12410–12415 (2019).

38. F. Alfei, K. Kanev, M. Hofmann, M. Wu, H. E. Ghoneim, P. Roelli, D. T. Utzschneider, M. von Hoesslin, J. G. Cullen, Y. Fan, V. Eisenberg, D. Wohlleber, K. Steiger, D. Merkler, M. Delorenzi, P. A. Knolle, C. J. Cohen, R. Thimme, B. Youngblood, D. Zehn, TOX reinforces the phenotype and longevity of exhausted T cells in chronic viral infection. Nature 571, 265–269 (2019).

39. A. C. Scott, F. Dundar, P. Zumbo, S. S. Chandran, C. A. Klebanoff, M. Shakiba, P. Trivedi, K. Menocal, H. Appleby, S. Camara, D. Zamarin, T. Walther, A. Snyder, M. R. Femia, E. A. Comen, H. Y. Wen, M. D. Hellmann, N. Anandasabapathy, Y. Liu, N. K. Altorki, P. Lauer, O. Levy, M. S. Glickman, J. Kaye, D. Betel, M. Philip, A. Schietinger, TOX is a critical regulator of tumour-specific T cell differentiation. Nature 571, 270–274 (2019).

40. C. Yao, H. W. Sun, N. E. Lacey, Y. Ji, E. A. Moseman, H. Y. Shih, E. F. Heuston, M. Kirby, S. Anderson, J. Cheng, O. Khan, R. Handon, J. Reilley, J. Fioravanti, J. Hu, S. Gossa, E. J. Wherry, L. Gattinoni, D. B. McGavern, J. J. O’Shea, P. L. Schwartzberg, T. Wu, Single-cell RNA-seq reveals TOX as a key regulator of CD8(+) T cell persistence in chronic infection. Nat Immunol 20, 890–901 (2019).

41. X. Wang, Q. He, H. Shen, A. Xia, W. Tian, W. Yu, B. Sun, TOX promotes the exhaustion of antitumor CD8(+) T cells by preventing PD1 degradation in hepatocellular carcinoma. J Hepatol 71, 731–741 (2019).

42. W. H. Hudson, J. Gensheimer, M. Hashimoto, A. Wieland, R. M. Valanparambil, P. Li, J. X. Lin, B. T. Konieczny, S. J. Im, G. J. Freeman, W. J. Leonard, H. T. Kissick, R. Ahmed, Proliferating Transitory T Cells with an Effector-like Transcriptional Signature Emerge from PD-1(+) Stem-like CD8(+) T Cells during Chronic Infection. Immunity 51, 1043–1058 e1044 (2019).

43. R. Zander, D. Schauder, G. Xin, C. Nguyen, X. Wu, A. Zajac, W. Cui, CD4(+) T Cell Help Is Required for the Formation of a Cytolytic CD8(+) T Cell Subset that Protects against Chronic Infection and Cancer. Immunity 51, 1028–1042 e1024 (2019).

44. Y. Chen, R. A. Zander, X. Wu, D. M. Schauder, M. Y. Kasmani, J. Shen, S. Zheng, R. Burns, E. J. Taparowsky, W. Cui, BATF regulates progenitor to cytolytic effector CD8(+) T cell transition during chronic viral infection. Nat Immunol 22, 996–1007 (2021).

45. K. S. Rome, S. J. Stein, M. Kurachi, J. Petrovic, G. W. Schwartz, E. A. Mack, S. Uljon, W. W. Wu, A. G. DeHart, S. E. McClory, L. Xu, P. A. Gimotty, S. C. Blacklow, R. B. Faryabi, E. J. Wherry, M. S. Jordan, W. S. Pear, Trib1 regulates T cell differentiation during chronic infection by restraining the effector program. J Exp Med 217, (2020).

46. S. E. McClory, O. Bardhan, K. S. Rome, J. R. Giles, A. E. Baxter, L. Xu, P. A. Gimotty, R. B. Faryabi, E. J. Wherry, W. S. Pear, M. S. Jordan, The pseudokinase Trib1 regulates the transition of exhausted T cells to a KLR(+) CD8(+) effector state, and its deletion improves checkpoint blockade. Cell Rep 42, 112905 (2023).

47. C. X. Dominguez, R. A. Amezquita, T. Guan, H. D. Marshall, N. S. Joshi, S. H. Kleinstein, S. M. Kaech, The transcription factors ZEB2 and T-bet cooperate to program cytotoxic T cell terminal differentiation in response to LCMV viral infection. J Exp Med 212, 2041–2056 (2015).

48. T. Guan, C. X. Dominguez, R. A. Amezquita, B. J. Laidlaw, J. Cheng, J. Henao-Mejia, A. Williams, R. A. Flavell, J. Lu, S. M. Kaech, ZEB1, ZEB2, and the miR-200 family form a counterregulatory network to regulate CD8(+) T cell fates. J Exp Med 215, 1153–1168 (2018).

49. C. Y. Yang, J. A. Best, J. Knell, E. Yang, A. D. Sheridan, A. K. Jesionek, H. S. Li, R. R. Rivera, K. C. Lind, L. M. D’Cruz, S. S. Watowich, C. Murre, A. W. Goldrath, The transcriptional regulators Id2 and Id3 control the formation of distinct memory CD8+ T cell subsets. Nat Immunol 12, 1221–1229 (2011).

50. D. Herndler-Brandstetter, H. Ishigame, R. Shinnakasu, V. Plajer, C. Stecher, J. Zhao, M. Lietzenmayer, L. Kroehling, A. Takumi, K. Kometani, T. Inoue, Y. Kluger, S. M. Kaech, T. Kurosaki, T. Okada, R. A. Flavell, KLRG1(+) Effector CD8(+) T Cells Lose KLRG1, Differentiate into All Memory T Cell Lineages, and Convey Enhanced Protective Immunity. Immunity 48, 716–729 e718 (2018).

51. K. R. Renkema, M. A. Huggins, H. Borges da Silva, T. P. Knutson, C. M. Henzler, S. E. Hamilton, KLRG1(+) Memory CD8 T Cells Combine Properties of Short-Lived Effectors and Long-Lived Memory. J Immunol 205, 1059–1069 (2020).

52. M. A. Turman, T. Yabe, C. McSherry, F. H. Bach, J. P. Houchins, Characterization of a novel gene (NKG7) on human chromosome 19 that is expressed in natural killer cells and T cells. Hum Immunol 36, 34–40 (1993).

53. X. Y. Li, D. Corvino, B. Nowlan, A. R. Aguilera, S. S. Ng, M. Braun, A. R. Cillo, T. Bald, K. J. Smyth, C. R. Engwerda, NKG7 Is Required for Optimal Antitumor T-cell Immunity. Cancer Immunol Res 10, 154–161 (2022).

54. S. Sarkar, V. Kalia, W. N. Haining, B. T. Konieczny, S. Subramaniam, R. Ahmed, Functional and genomic profiling of effector CD8 T cell subsets with distinct memory fates. J Exp Med 205, 625–640 (2008).

55. Z. Chen, Z. Ji, S. F. Ngiow, S. Manne, Z. Cai, A. C. Huang, J. Johnson, R. P. Staupe, B. Bengsch, C. Xu, S. Yu, M. Kurachi, R. S. Herati, L. A. Vella, A. E. Baxter, J. E. Wu, O. Khan, J. C. Beltra, J. R. Giles, E. Stelekati, L. M. McLane, C. W. Lau, X. Yang, S. L. Berger, G. Vahedi, H. Ji, E. J. Wherry, TCF-1-Centered Transcriptional Network Drives an Effector versus Exhausted CD8 T Cell-Fate Decision. Immunity 51, 840–855 e845 (2019).

56. J. R. Giles, S. Manne, E. Freilich, D. A. Oldridge, A. E. Baxter, S. George, Z. Chen, H. Huang, L. Chilukuri, M. Carberry, L. Giles, N. P. Weng, R. M. Young, C. H. June, L. M. Schuchter, R. K. Amaravadi, X. Xu, G. C. Karakousis, T. C. Mitchell, A. C. Huang, J. Shi, E. J. Wherry, Human epigenetic and transcriptional T cell differentiation atlas for identifying functional T cell-specific enhancers. Immunity 55, 557–574 e557 (2022).

57. J. Eyquem, J. Mansilla-Soto, T. Giavridis, S. J. van der Stegen, M. Hamieh, K. M. Cunanan, A. Odak, M. Gonen, M. Sadelain, Targeting a CAR to the TRAC locus with CRISPR/Cas9 enhances tumour rejection. Nature 543, 113–117 (2017).

58. S. M. Collins, K. A. Alexander, S. Lundh, A. J. Dimitri, Z. Zhang, C. R. Good, J. A. Fraietta, S. L. Berger, TOX2 coordinates with TET2 to positively regulate central memory differentiation in human CAR T cells. Sci Adv 9, eadh2605 (2023).

59. R. Saleh, V. Sasidharan Nair, S. M. Toor, R. Z. Taha, K. Murshed, M. Al-Dhaheri, M. Khawar, M. A. Petkar, M. Abu Nada, F. Al-Ejeh, E. Elkord, Differential gene expression of tumor-infiltrating CD8(+) T cells in advanced versus early-stage colorectal cancer and identification of a gene signature of poor prognosis. J Immunother Cancer 8, (2020).

60. W. Zheng, J. Wei, C. C. Zebley, L. L. Jones, Y. Dhungana, Y. D. Wang, J. Mavuluri, L. Long, Y. Fan, B. Youngblood, H. Chi, T. L. Geiger, Regnase-1 suppresses TCF-1+ precursor exhausted T-cell formation to limit CAR-T-cell responses against ALL. Blood 138, 122–135 (2021).

61. B. Zangari, T. Tsuji, J. Matsuzaki, H. Mohammadpour, C. Eppolito, S. Battaglia, F. Ito, T. Chodon, R. Koya, A. J. Robert McGray, K. Odunsi, Tcf-1 protects anti-tumor TCR-engineered CD8(+) T-cells from GzmB mediated self-destruction. Cancer Immunol Immunother 71, 2881–2898 (2022).

62. C. Tsui, L. Kretschmer, S. Rapelius, S. S. Gabriel, D. Chisanga, K. Knopper, D. T. Utzschneider, S. Nussing, Y. Liao, T. Mason, S. V. Torres, S. A. Wilcox, K. Kanev, S. Jarosch, J. Leube, S. L. Nutt, D. Zehn, I. A. Parish, W. Kastenmuller, W. Shi, V. R. Buchholz, A. Kallies, MYB orchestrates T cell exhaustion and response to checkpoint inhibition. Nature 609, 354–360 (2022).

63. G. Lopez-Cantillo, C. Uruena, B. A. Camacho, C. Ramirez-Segura, CAR-T Cell Performance: How to Improve Their Persistence? Front Immunol 13, 878209 (2022).

64. M. D. Buck, D. O’Sullivan, R. I. Klein Geltink, J. D. Curtis, C. H. Chang, D. E. Sanin, J. Qiu, O. Kretz, D. Braas, G. J. van der Windt, Q. Chen, S. C. Huang, C. M. O’Neill, B. T. Edelson, E. J. Pearce, H. Sesaki, T. B. Huber, A. S. Rambold, E. L. Pearce, Mitochondrial Dynamics Controls T Cell Fate through Metabolic Programming. Cell 166, 63–76 (2016).

65. L. Gattinoni, E. Lugli, Y. Ji, Z. Pos, C. M. Paulos, M. F. Quigley, J. R. Almeida, E. Gostick, Z. Yu, C. Carpenito, E. Wang, D. C. Douek, D. A. Price, C. H. June, F. M. Marincola, M. Roederer, L. P. Restifo, A human memory T cell subset with stem cell-like properties. Nat Med 17, 1290–1297 (2011).

66. X. Wang, L. L. Popplewell, J. R. Wagner, A. Naranjo, M. S. Blanchard, M. R. Mott, A. P. Norris, C. W. Wong, R. Z. Urak, W. C. Chang, S. K. Khaled, T. Siddiqi, L. E. Budde, J. Xu, B. Chang, N. Gidwaney, S. H. Thomas, L. J. Cooper, S. R. Riddell, …, S. J. Forman, Phase 1 studies of central memory-derived CD19 CAR T-cell therapy following autologous HSCT in patients with B-cell NHL. Blood 127, 2980–2990 (2016).

67. L. Biasco, N. Izotova, C. Rivat, S. Ghorashian, R. Richardson, A. Guvenel, R. Hough, R. Wynn, B. Popova, A. Lopes, M. Pule, A. J. Thrasher, P. J. Amrolia, Clonal expansion of T memory stem cells determines early anti-leukemic responses and long-term CAR T cell persistence in patients. Nat Cancer 2, 629–642 (2021).

68. N. E. Scharping, A. V. Menk, R. S. Moreci, R. D. Whetstone, R. E. Dadey, S. C. Watkins, R. L. Ferris, G. M. Delgoffe, The Tumor Microenvironment Represses T Cell Mitochondrial Biogenesis to Drive Intratumoral T Cell Metabolic Insufficiency and Dysfunction. Immunity 45, 374–388 (2016).

69. H. Kunimoto, H. Nakajima, TET2: A cornerstone in normal and malignant hematopoiesis. Cancer Sci 112, 31–40 (2021).

70. S. Mustjoki, N. S. Young, Somatic Mutations in “Benign” Disease. N Engl J Med 384, 2039–2052 (2021).

71. E. A. Stadtmauer, J. A. Fraietta, M. M. Davis, A. D. Cohen, K. L. Weber, E. Lancaster, P. A. Mangan, I. Kulikovskaya, M. Gupta, F. Chen, L. Tian, V. E. Gonzalez, J. Xu, I. Y. Jung, J. J. Melenhorst, G. Plesa, J. Shea, T. Matlawski, A. Cervini, A. L. Gaymon, S. Desjardins, A. Lamontagne, J. Salas-Mckee, A. Fesnak, D. L. Siegel, B. L. Levine, J. K. Jadlowsky, R. M. Young, A. Chew, W. T. Hwang, E. O. Hexner, B. M. Carreno, C. L. Nobles, F. D. Bushman, K. R. Parker, Y. Qi, A. T. Satpathy, H. Y. Chang, Y. Zhao, S. F. Lacey, C. H. June, CRISPR-engineered T cells in patients with refractory cancer. Science 367, (2020).

72. L. Couronne, C. Bastard, O. A. Bernard, TET2 and DNMT3A mutations in human T-cell lymphoma. N Engl J Med 366, 95–96 (2012).

73. X. Zhang, J. Su, M. Jeong, M. Ko, Y. Huang, H. J. Park, A. Guzman, Y. Lei, Y. H. Huang, A. Rao, W. Li, M. A. Goodell, DNMT3A and TET2 compete and cooperate to repress lineage-specific transcription factors in hematopoietic stem cells. Nat Genet 48, 1014–1023 (2016).

74. J. Garcia, J. Daniels, Y. Lee, I. Zhu, K. Cheng, Q. Liu, D. Goodman, C. Burnett, C. Law, C. Thienpont, J. Alavi, C. Azimi, G. Montgomery, K. T. Roybal, J. Choi, Naturally occurring T cell mutations enhance engineered T cell therapies. Nature, (2024).

75. C. L. Nobles, S. Sherrill-Mix, J. K. Everett, S. Reddy, J. A. Fraietta, D. L. Porter, N. Frey, S. I. Gill, S. A. Grupp, S. L. Maude, D. L. Siegel, B. L. Levine, C. H. June, S. F. Lacey, J. J. Melenhorst, F. D. Bushman, CD19-targeting CAR T cell immunotherapy outcomes correlate with genomic modification by vector integration. J Clin Invest 130, 673–685 (2020).

76. E. Sherman, C. Nobles, C. C. Berry, E. Six, Y. Wu, A. Dryga, N. Malani, F. Male, S. Reddy, A. Bailey, K. Bittinger, J. K. Everett, L. Caccavelli, M. J. Drake, P. Bates, S. Hacein-Bey-Abina, M. Cavazzana, F. D. Bushman, INSPIIRED: A Pipeline for Quantitative Analysis of Sites of New DNA Integration in Cellular Genomes. Molecular Therapy - Methods & Clinical Development 4, 39–49 (2017).

77. W. Kong, A. Dimitri, W. Wang, I. Y. Jung, C. J. Ott, M. Fasolino, Y. Wang, I. Kulikovskaya, M. Gupta, T. Yoder, J. E. DeNizio, J. K. Everett, E. F. Williams, J. Xu, J. Scholler, T. J. Reich, V. G. Bhoj, K. M. Haines, M. V. Maus, …, J. A. Fraietta, BET bromodomain protein inhibition reverses chimeric antigen receptor extinction and reinvigorates exhausted T cells in chronic lymphocytic leukemia. J Clin Invest 131, (2021).

78. H. Pircher, J. Baenziger, M. Schilham, T. Sado, H. Kamisaku, H. Hengartner, R. M. Zinkernagel, Characterization of virus-specific cytotoxic T cell clones from allogeneic bone marrow chimeras. Eur J Immunol 17, 159–166 (1987).

79. H. Pircher, E. E. Michalopoulos, A. Iwamoto, P. S. Ohashi, J. Baenziger, H. Hengartner, R. M. Zinkernagel, T. W. Mak, Molecular analysis of the antigen receptor of virus-specific cytotoxic T cells and identification of a new V alpha family. Eur J Immunol 17, 1843–1846 (1987).

80. P. M. Odorizzi, K. E. Pauken, M. A. Paley, A. Sharpe, E. J. Wherry, Genetic absence of PD-1 promotes accumulation of terminally differentiated exhausted CD8+ T cells. J Exp Med 212, 1125–1137 (2015).

81. M. Kurachi, J. Kurachi, Z. Chen, J. Johnson, O. Khan, B. Bengsch, E. Stelekati, J. Attanasio, L. M. McLane, M. Tomura, S. Ueha, E. J. Wherry, Optimized retroviral transduction of mouse T cells for in vivo assessment of gene function. Nat Protoc 12, 1980–1998 (2017).

82. A. E. Baxter, H. Huang, J. R. Giles, Z. Chen, J. E. Wu, S. Drury, K. Dalton, S. L. Park, L. Torres, B. W. Simone, M. Klapholz, S. F. Ngiow, E. Freilich, S. Manne, V. Alcalde, V. Ekshyyan, S. L. Berger, J. Shi, M. S. Jordan, E. J. Wherry, The SWI/SNF chromatin remodeling complexes BAF and PBAF differentially regulate epigenetic transitions in exhausted CD8(+) T cells. Immunity 56, 1320–1340 e1310 (2023).

83. J. D. Buenrostro, P. G. Giresi, L. C. Zaba, H. Y. Chang, W. J. Greenleaf, Transposition of native chromatin for fast and sensitive epigenomic profiling of open chromatin, DNA-binding proteins and nucleosome position. Nat Methods 10, 1213–1218 (2013).

